# To Kill or not to Kill: A Conserved *trans*-intoxication protection factor Blocks X-T4SS-Mediated Fratricide through Interaction with VirB5

**DOI:** 10.64898/2026.05.09.723464

**Authors:** Gabriel U. Oka, Camilla Adan, Thiago Rodrigo dos Santos, William Cenens, Diorge P. Souza, Chuck S. Farah

**Author notes:** Instituto de Física de São Carlos, Universidade de São Paulo, Avenida João Dagnone, n° 1100, São Carlos, CEP 13563-120, Brazil.

## Abstract

Bacteria can outcompete rivals by using specialized secretion systems to deliver toxic effectors into prey cells. How these systems distinguish prey from kin remains a fundamental unanswered question. In many Xanthomonadales (Lysobacterales) species, the antibacterial type IV secretion system (X-T4SS) mediates cell–cell contact-dependent killing of competing bacteria. Here, we identified XAC2611, a chromosomally encoded, cysteine-rich DUF4189 protein, as essential for preventing X-T4SS-mediated fratricide among sibling cells in *Xanthomonas citri*. XAC2611 homologs are found within or proximal to the genomic loci coding almost all identified X-T4SS. Live-cell microscopy and biochemical data reveal that XAC2611 is abundantly produced and widely distributed throughout the periplasm of recipient cells and is required to prevent *X. citri* cells from intoxicating each other in a contact-dependent manner. We also show that XAC2611 homologs from *Stenotrophomonas maltophilia* protect *X. citri* cells from attack by *S. maltophilia.* Protein-protein interaction assays show that XAC2611 interacts directly with the VirB5 subunit. Since VirB5 is predicted to be localized at the tip of the X-T4SS pilus, its interaction with XAC2611 in a neighboring sister cell could block X-T4SS-mediated effector delivery. We therefore name this family of proteins *trans*-intoxication protection factors (Tpfs).

## INTRODUCTION

Bacteria colonize a wide range of structured and dynamic environments, including soil, the rhizosphere, aquatic ecosystems, biofilms, the animal oral cavity and digestive tract, where competition for nutrients and space shapes microbial evolution and ecological diversification (Hibbing et al. 2010). In these environments, interbacterial competition through the elimination of rival bacteria is widespread and frequently mediated by specialized bacterial secretion systems (Granato et al. 2019). Among these, a chromosomally encoded Type IV Secretion System (X-T4SS), broadly distributed across species of the order *Lysobacterales (earlier known as Xanthomonadales (Kumar et al. 2019))* (X-T4SS), plays a central role in killing competing bacteria (Souza et al. 2015; Sgro et al. 2019; Bayer-Santos et al. 2019; Shen et al. 2021; Cobe et al. 2024; Oka et al. 2022; Alegria et al. 2005)).

The X-T4SS machinery is structurally similar to the canonical type IVA secretion system, comprising 12 conserved VirB proteins (VirB1–VirB11) and the coupling protein VirD4 (Alvarez-Martinez and Christie 2009; Sgro et al. 2019). In *Xanthomonas citri*, the X-T4SS translocates a family of polymorphic antibacterial effectors (X-Tfes) characterized by diverse N-terminal toxic domains and a conserved C-terminal XVIPCD domain, which mediates recruitment to the secretion machinery through direct interaction with the all-alpha domain of the coupling protein VirD4 (Alegria et al. 2005; Souza et al. 2015; Oka et al. 2022). Once recruited, X-Tfes are secreted into neighboring target cells in a contact-dependent manner, causing rapid lysis (Souza et al. 2015; Oka et al. 2022). To prevent self-intoxication, X-Tfes are usually encoded together with their corresponding immunity proteins (X-Tfis) in bicistronic operons (Souza et al. 2015; Oka et al. 2022, 2024; Bayer-Santos et al. 2019; Sgro et al. 2019).

X-Tfis protect against *cis*-intoxication, an intracellular process exemplified by the effector X-Tfe^XAC2609^, which is produced in the cytoplasm and causes damage in the periplasm when its cognate immunity protein, X-Tfi^XAC2610^, is absent (Oka et al. 2024). Notably, this toxicity occurs independently of a functional X-T4SS, highlighting that X-Tfis are essential for preventing intracellular self- or *cis*-intoxication (Oka et al. 2024). In several other antibacterial systems, including the T1SS (García-Bayona et al. 2017), T5SS (Aoki et al. 2005), T6SS (Russell et al. 2011), and the T7SSb of Gram-positive bacteria (Cao et al. 2016; Tassinari et al. 2022; Kobayashi 2021), protection against *trans*-intoxication among sibling cells has traditionally been explained by the paradigmatic model of cognate toxin–immunity protein pairs. However, *X. citri* wild-type strains fail to kill *X. citri* mutants lacking up to eight X-Tfe/X-Tfi pairs (Oka et al. 2024), indicating that X-Tfis are not the primary line of defense against effectors delivered by neighboring sibling cells. Thus, it remains unclear whether bacteria carrying an X-T4SS possess a dedicated kin-recognition mechanism to prevent *trans*-intoxication among genetically identical cells.

Here, we investigate the molecular basis of *trans*-intoxication prevention mediated by the X-T4SS in *Xanthomonas citri*. Through the use of knockout strains, interbacterial competition assays, live-cell fluorescence microscopy, and biochemical analysis, we identify XAC2611, a cysteine-rich DUF4189 protein encoded in the X-T4SS gene cluster, as essential for protecting sibling cells from X-T4SS-mediated fratricide. Our data show that XAC2611 is abundantly produced and localized in the periplasm of recipient cells, where it interacts directly with the VirB5 subunit of the X-T4SS which is expected to be localized at the tip of the emerging pilus of a genetically identical donor (Macé and Waksman 2024). Genes coding for XAC2611 homologs can be found associated with all the *loci* coding for X-T4SS and we show that the heterologous expression of a one or more *Stenotrophomonas maltophilia* XAC2611 homolog can protect *X. citri* from X-T4SS-dependent attack by *S. maltophilia*. This mechanism represents a previously unrecognized, species-specific form of kin recognition, enabling X-T4SS-bearing bacteria to selectively block effector delivery into sibling cells and presents interesting parallels with previously described surface and entry exclusion mechanisms in conjugative T4SSs.

## RESULTS

### A conserved DUF4189 protein found associated with X-T4SSs

In *X. citri*, the genes coding for the components of the X-T4SS are arranged collinearly in the *vir* locus (Alegria et al. 2005; Sgro et al. 2019). In addition to the structural genes for the components of the X-T4SS (VirD4, VirB1-VirB11), the *X. citri vir* locus also codes for an effector/immunity protein pair (X-Tfe^XAC2609^/X-Tfi^XAC2610^) and two homologous proteins XAC2611 and XAC2606. The intervening open reading frames (ORFs) XAC2607 and XAC2608 encode truncated versions of VirB6. Interestingly, *xac2611*, but not *xac2606*, is part of a polycistronic transcript that extends from *virB7* to *xac2609* (Cenens et al. 2020). XAC2611 and XAC2606 contain a domain of unknown function (InterPro: IPR025240, DUF4189) preceded by a putative cleavable N-terminal signal sequence (**Figure 1**), suggestive of their localization in the bacterial cell envelope. Members of this homologous superfamily are predicted by Alphafold3 (Abramson et al. 2024), to adopt a conserved topology stabilized by three highly conserved disulfide bridges (**Figure 1C**). Genes coding for homologs of XAC2611 and XAC2606 can be found within or in the immediate vicinity of *loci* that code for X-T4SS in many other genera, including *Stenotrophomonas spp*, *Lysobacter spp, Luteibacter ssp and Dyella ssp* (Figure **Figure 1**; (Alegria et al. 2005; Sgro et al. 2019)). The genes are most commonly immediately downstream of the VirB6 gene and in many cases are unannotated and occur in multiple copies. The presence of XAC2611 and XAC2606 homologues across multiple antibacterial X-T4SS gene clusters and their predicted cellular location point to them as possible candidates to inhibit *trans*-intoxication in *Lysobacterales* species.

**Figure 1.**
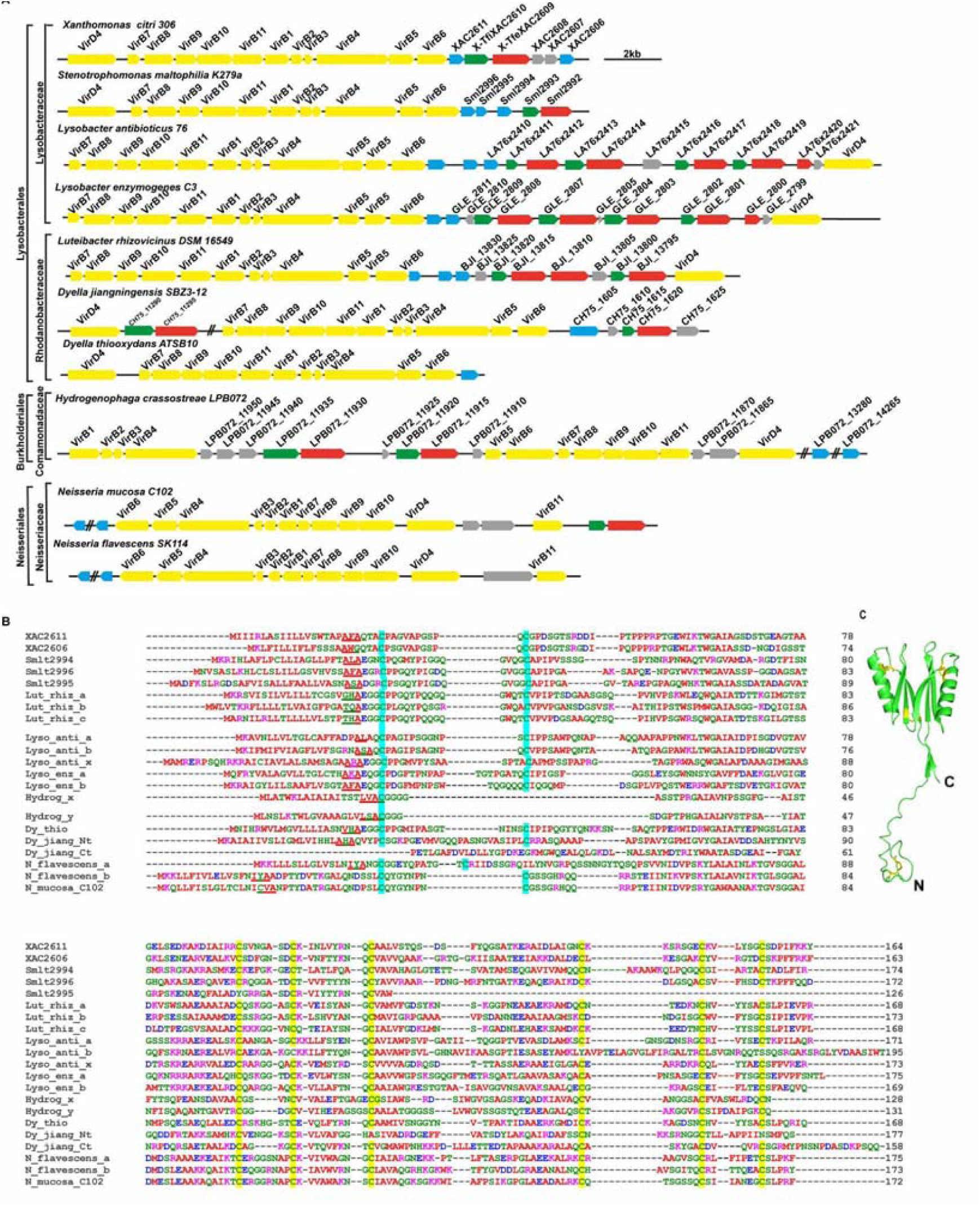
**A)** *Loci* coding for X-T4SS in representative bacterial species. XAC2611 homologs are indicated in cyan. Yellow, red, green and grey boxes indicate genes that code for putative T4SS channel components, effectors, immunity proteins and genes of unknown function, respectively. In several cases, the homologs were not annotated in the deposited genomes. **B)** Alignment of XAC2611 and homologs found in the genomes of representative bacteria coding for X-T4SSs. The six conserved cysteine residues found in the globular C-terminal domain are highlighted in yellow and the two conserved cysteine residues found in the predicted intrinsically disordered N-terminal domain are highlighted in cyan. The N-terminal signal peptides are shown in bold type with signal peptidase I (SPI) cleavage (AxA) motif underlined (Auclair et al. 2012). Sec signal peptide Signal Peptidase I cleavage site were identified by SignalP 6.0 (Teufel et al. 2022). Bacterial species: *X. citri* 306 (XAC2611, XAC2606); *S. maltophilia* K279 (Smlt2994, Smlt 2995, Smlt2996); *Luteibacter rhizovicinus* DSM16549 (Lut_rhiz_a, Lut_rhiz_b, Lut_rhiz_c); *Lysobacter antibioticus* 76 (Lyso_anti_a, Lyso_anti_b, Lyso_anti_x); *Lysobacter enzymogenes* C3 (Lyso_enz_a, Lysoenz_b); *Hydrogenophaga crassostreae* strain LPB0072 (Hydrogen_x, Hydrogen_y); Dyella thiooxydans ATSB10 (Dy_thio); *Dyella jiangningensis* JB7_3-12 N-terminal and C-terminal domains (Dy_jiang_Nt, Dy_jiang_Ct, respectively); *Neisseria flavescens* SK114 (N_flavescens_a, N-flavescens_b; *Neisseria mucosa* C102 (N_mucosa_C102). Inset: **C)** Ribbon representation of the Alphafold model of XAC2611 (residues 22-164). Disulphide bridges are shown in yellow.

### XAC2611 is localized in the *X. citri* cell envelope

We used fluorescence microscopy to experimentally detect the expression and localization of XAC2611 and XAC2606 in *X. citri* cells expressing the proteins as msfGFP chimeras (**Figure 2A**). We observed that XAC2611-GFP is highly expressed and localized to the cell envelope. In contrast, we could not detect any significant expression of XAC2606-GFP under these growth conditions (**Figure 2A**). We therefore decided to focus on XAC2611 and aimed to determine whether it is exposed on the outside of the cell membrane or localized within the cell envelope. To address this, we fixed live *X. citri* carrying a plasmid expressing XAC2611 with a C-terminal FLAG tag (pBRA-XAC2611-FLAG cells) with formaldehyde, followed by washing with either water or 70% ethanol, the latter of which permeabilizes the cell envelope (Park et al. 2019; Fei et al. 2015). The cells were then incubated with or without lysozyme to increase access to periplasmic proteins (**Figure 2B**). To detect exposed XAC2611-FLAG we used antiFLAG antibodies with a secondary anti-IgG antibody conjugated with Alexafluor. **Figure 2C** shows that XAC2611-FLAG was only detected after treatment with ethanol and lysozyme, suggesting that the protein is not exposed on the external surface of the cell but rather within the cell envelope. These results are consistent with a recent study that detected XAC2611 in *X. citri* outer membrane vesicles (Araujo et al. 2025). Periplasmatic localization is also consistent with the observation that XAC2611 and its orthologs all have a predicted N-terminal signal peptide with a signal peptidase I (SPI) cleavage (AxA) motif (**Figure 1B**). Interestingly, the predicted SPI cleavage site is followed by a highly conserved cysteine residue (**Figure 1B**). Cysteine residues associated with signal peptides are often contained within lipobox motifs (consensus sequence [LVI][ASTVI][GAS]C at positions -3, -2, -1, +1) which mediate cleavage and lipidation at the invariant N-terminal cysteine residue (Hayashi and Wu 1990; Babu et al. 2006). However, in the case of XAC2611 and its homologs, the lipobox consensus sequence is not present (**Figure 1B**) suggesting that the protein may be transported and processed (via SP1 cleavage at the AxA motif) as soluble components within the periplasm.

**Figure 2.**
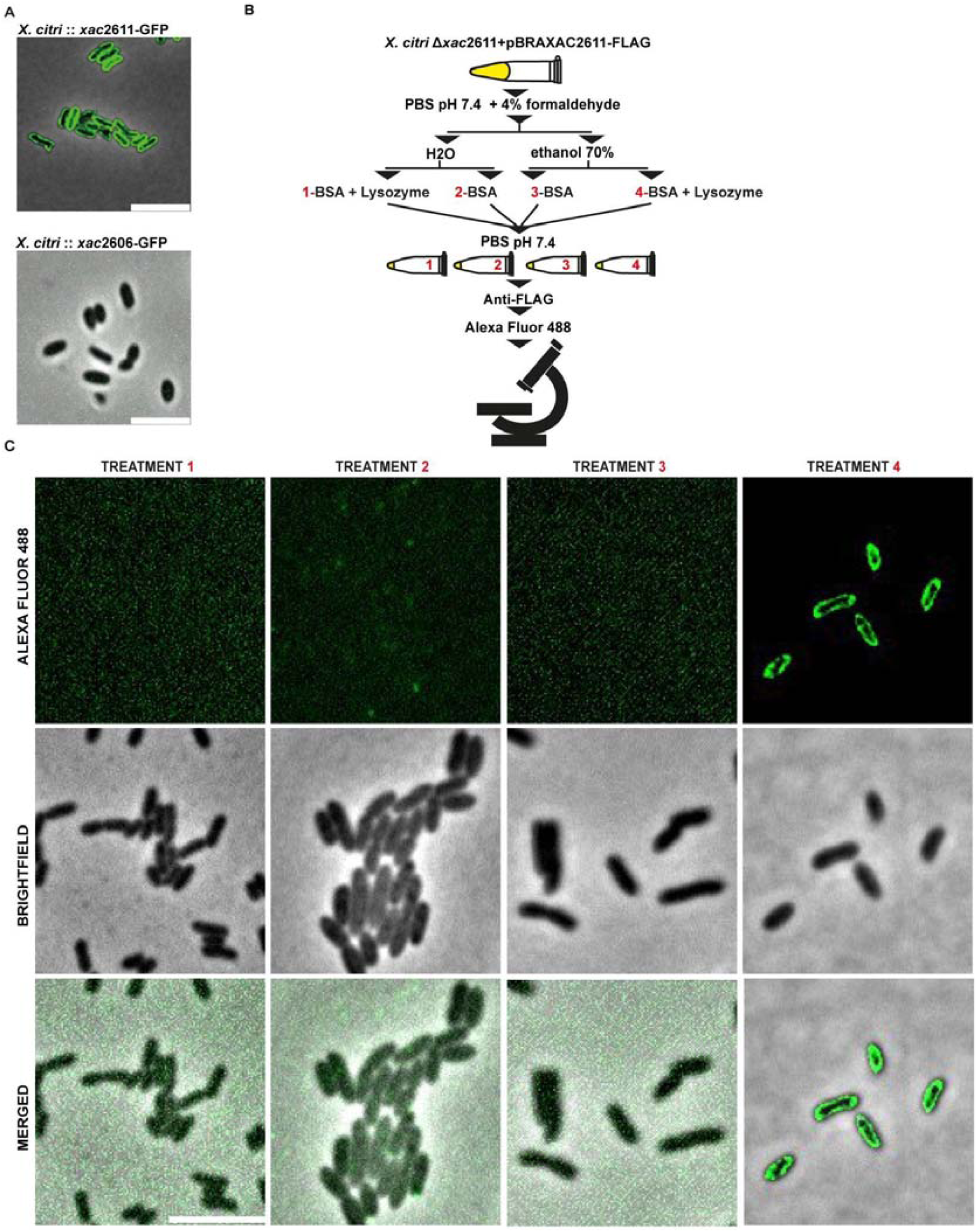
XAC2611 is localized in the interior of the bacterial envelope. **(A)** Fluorescence microscopy images of *X. citri* cells in which msfGFP genes were fused to the 3’ end of *xac2611* (*xac*2611-GFP) or *xac2606* (*xac*2606-GFP). The image shows the msfGFP intensity at the focal plane of the cells. Scale bars, 5 μm. **(B)** Schematic of procedure for permeabilization of *X. citri* Δ*xac*2611 cells transformed with the pBRA-XAC2611-FLAG plasmid. Briefly, *X. citri* cells were grown in 2xTY medium and then fixed in PBS solution supplemented with 4% formaldehyde in microtubes. Subsequently, four distinct treatments were performed: (1) Incubation in water followed by incubation in PBS supplemented with 1% BSA; (2) Incubation in water followed by incubation in PBS supplemented with 1% BSA and lysozyme; (3) Incubation in a 70% ethanol solution followed by incubation in PBS supplemented with 1% BSA; (4) Incubation in a 70% ethanol solution followed by incubation in PBS supplemented with 1% BSA and lysozyme. Following these treatments, samples were incubated with Anti-FLAG and Anti-IgG antibodies conjugated with Alexa Fluor® 488 (ab150077). See material and methods for details. (**C)** *In situ detection* of XAC2611-FLAG. Following the treatments as outlined in (C), cells presenting Alexa Fluor®488 fluorescence were visualized by microscopy. Images show a blind deconvoluted (for treatment 4) slice of the central plane of the fluorescence channel (Alexa fluor 488), Phase Contrast (Brightfield), and the merged images of both channels. Scale bar 5 μm.

### XAC2611 is required to avoid X-T4SS-dependent lysis in *X. citri*

We then set out to explore the *in vivo* function of the 164 residue XAC2611 protein. **Supplementary Figures 1 and 2** show that X-T4SS-mediated killing of *E. coli* is maintained in the Δ*xac*2611 mutant but is not observed for the *X. citri* Δ*virB7 and X. citri* Δ*virB5* mutants, thus demonstrating that the XAC2611 is not required for X-T4SS function. We then generated in-frame knockouts of the *xac2611* gene in an *X. citri* strain carrying a plasmid expressing red fluorescent protein (RFP), for visualization using fluorescence time-lapse microscopy. **Figure 3 and Movies S1-S6** show time-lapse images of these *X. citri* strains grown over a 5 hour period on agarose slabs. Wild-type *X. citri* cells grow normally over this time period as reflected by the constant RFP fluorescence (**Figure 3A** and **Movie S1**). However, during the growth of Δ*xac*2611 cells, many cells lost their fluorescence, often suffering significant reductions in size and loss of morphology indicative of lysis. After 5 hours, approximately 40% of Δ*xac*2611 cells exhibited signs of cell death (**Figure 3B** and **Movie S2**).

**Figure 3.**
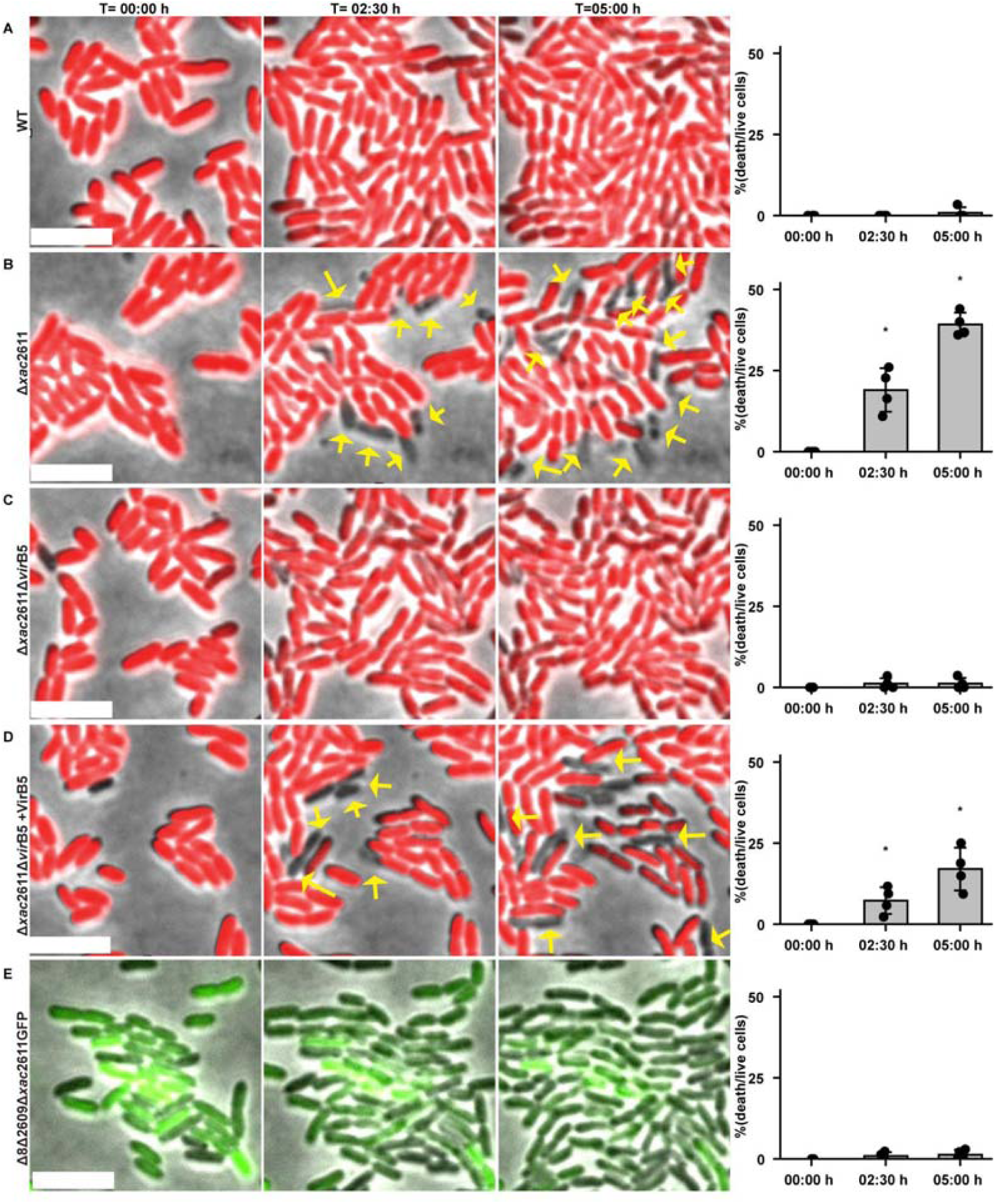
The absence of XAC2611 induces X-T4SS-dependent lysis in *X. citri* monocultures. **(A-E)** Time-lapse microscopy images depict representative fields shown in movie S1 at the onset of the experiment at (T=00:00 h), 2.5 hours in (T=02:30 h), and at 5 hours (05:00 h). Cellular lysis events are highlighted with arrows. *X. citri* strains employed in the assay all carry a plasmid encoding for RFP or GFP. Wild type (WT), knockout of *xac2611* strain (Δ*xac*2611), double knockout of *xac2611* and *virB5* (Δ*xac*2611ΔvirB5), Δ8Δ2609Δ*xac*2611GFP and the double knockout strain Δ*xac*2611Δ*virB5* carrying the plasmid pBRA-VirB5 Δ*xac*2611Δ*vir*B5 + VirB5). Total cumulative deaths at T = 02:30 h were: WT, 0; Δ*xac*2611, 36;Δ*xac*2611ΔvirB5, 3; Δ*xac*2611Δ*vir*B5 + VirB5, 17; and Δ8Δ2609GFPΔ*xac*2611, 3. At T = 05:00 h, totals were: WT, 3; Δ*xac*2611, 89; Δ*xac*2611Δ*vir*B5, 4; Δ*xac*2611Δ*vir*B5 + VirB5, 59; and Δ8Δ2609GFPΔ*xac*2611, 5. Statistics were performed independently at each time point by one-way ANOVA followed by Dunnett’s test versus WT. At T = 02:30 h, adjusted p-values were: Δ*xac*2611, 7.26 × 10[[; Δ*xac*2611Δ*vir*B5, 0.973; Δ*xac*2611Δ*vir*B5 + VirB5, 0.0426; and Δ8Δ2609Δ*xac*2611GFP, 0.988. At T = 05:00 h, adjusted p-values were: Δxac2611, 7.77 × 10[¹[;Δ*xac*2611Δ*vir*B5, 0.9998; Δ*xac*2611Δ*vir*B5 + VirB5, 7.01 × 10[[; and Δ8Δ2609Δ*xac*2611GFP, 0.9993. Statistical significance was assessed using one-way ANOVA followed by Dunnett’s multiple-comparison test against WT, with α = 0.05. Scale bar, 5 μm.

### X-T4SS-dependent *trans*-intoxication is inhibited by XAC2611

We then asked whether the death of cells lacking XAC2611 is due to *trans*-intoxication, in other words, the X-T4SS dependent transfer of X-Tfe toxins from one *X. citri* cell to another. If this is the case, the lysis events observed for *X. citri* Δ*xac*2611 cells should not be observed if X-T4SS function were disrupted. **Figure 3C** and **Supplementary Movie 3** show that nearly no cell death is observed in monocultures of the Δ*xac*2611Δ*virB5* double mutant. However, cell death is recovered when we transform the Δ*xac*2611Δ*virB5* strain with a plasmid-directing expression of VirB5 (**Figure 3D** and **Supplementary Movie 4**). Therefore cell death is dependent on a functional X-T4SS. To test whether cell death is also dependent on the presence of X-Tfes, we deleted the *xac2611* gene in the *X. citri* Δ8Δ2609GFP strain which lacks 9 X-Tfes and has been shown to be unable to kill *E. coli* in interspecies competition experiments (Oka et al. 2022, 2024). **Figure 3E** and **Supplementary movie 5** show that in the absence of X-Tfes, the lack of XAC2611 does not lead to *trans*-intoxication.

To further confirm that the death of cells lacking XAC2611 is due to *trans*-intoxication, we used fluorescence microscopy to observe co-cultures of different combinations of *X. citri* wild type and mutant cells expressing GFP or RFP (**Figure 4 and Supplementary Movies 7-12)**. We have previously shown that the *X. citri* Δ8Δ2609 strain that lacks nine X-Tfes and eight X-Tfis is not killed by *X. citri* wild-type cells (Oka et al. 2024). This observation was confirmed using wild-type *X. citri*-GFP cells in co-culture with Δ8Δ2609 RFP cells (**Figure 4A, Supplementary movie 7**). However, *X. citri* WT cells are able to kill Δ8Δ2609Δ*xac*2611 cells (**Figure 4B, Supplementary Movie 8**). These observations are consistent with an X-T4SS-dependent *trans*-intoxication mechanism that is inhibited by XAC2611. **Figure 4C and Supplementary Movie 9** show micrographs of cocultures of the Δ8Δ2609GFP and Δ8Δ2609Δ*xac*2611RFP cells where, as expected, no death of the latter strain was observed. Additionally, deletion of the *xac2606* gene, that codes for a XAC2611 paralog, in the Δ8Δ2609 background did not lead to significant *trans*-intoxication by wild-type *X. citri* cells (**Supplementary Movie 10**), consistent with the hypothesis that XAC2611 provides the major contribution of protection against *trans*-intoxication under the conditions tested. Moreover, *trans*-intoxication mediated by the X-T4SS was confirmed in experiments using a Δ*virB5* knockout strain against the Δ8Δ2609Δ*xac*2611RFP recipient, where no killing was observed (**Figure 4D and Supplementary Movie 11**). Killing was restored upon episomal complementation of the Δ*virB5* strain (**Figure 4E and Supplementary Movie 12**). Finally, as expected, killing was abrogated by complementing the Δ8Δ2609Δ*xac*2611RFP recipient with episomal expression of XAC2611 (**Figure 4F and Supplementary Movie 13).**

**Figure 4.**
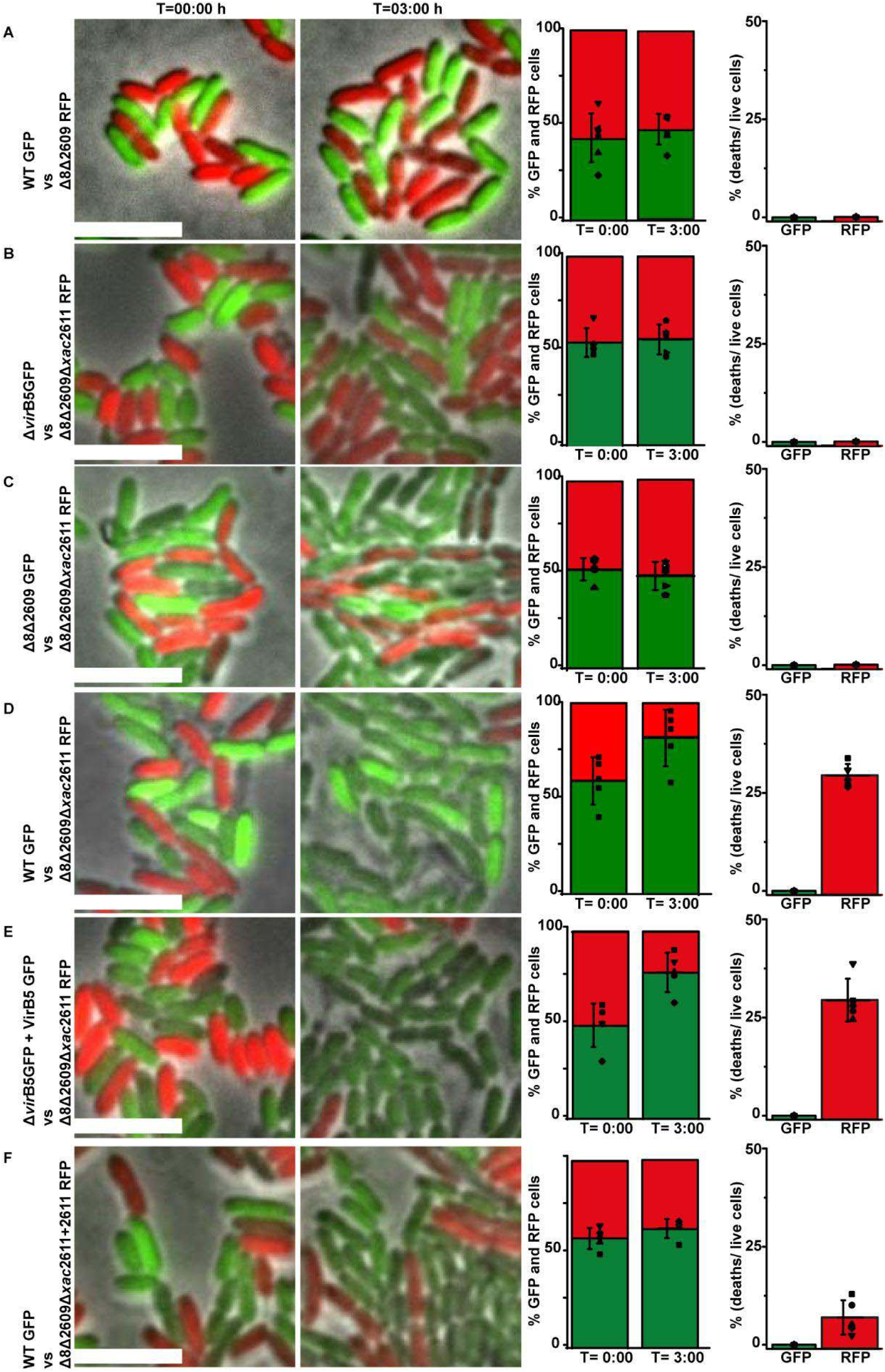
Fratricide in mixed *X. citri* co-cultures caused by knockout of *xac*2611. **(A-F)** Time-lapse fluorescence microscopy of a mixed culture of *X. citri* strains labeled with either GFP (green) or RFP (red) at the start of the experiment (T=00:00 h) and after 3 hours (T=03:00 h). *X. citri* strains include wild type (WT), Δ*vir*B5 (ΔVirB5), Δ8Δ2609, Δ8Δ2609 with *xac2611* deletion (Δ8Δ2609Δ*xac*2611), *X. citri* Δ8Δ2609Δ*xac*2611 transformed with plasmid pBRA-XAC2611, and *X. citri* Δ*vir*B5 + pBRA-VirB5. The left graph bars represent the mean percentage from visually identified GFP and RFP-tagged cells at the experiment’s onset (T=0:00) and after 3 hours (T=3:00), while the right graph bars depict the relative cumulative lysis events, that was calculated by counting the number of death events recorded during 3 hours for GFP or RFP tagged cells divided by the total number of cells counted at the start, observed across five to six representative microscopy fields shown in movies S7 to S12. Symbols denote counts for each field, bars indicate mean, and error bars represent standard deviation. Scale bar: 5 μm.

To quantify the extent to which XAC2611-dependent inhibition of *trans*-intoxication contributes to the vitality of an *X. citri* population, we performed co-culture competition assays in which equal numbers of cells of two *X. citri* strains carrying plasmids conferring kanamycin (pBBRGFP) or gentamicin (pBBRRFP) resistance were grown for 40 hours in 2xTY agar plates after which the number of colony forming units (CFUs) of each strain was quantified using selective media. **Figure 5** presents the results of these interbacterial competition experiments. *X. citri* wild-type exhibited a 230-fold viability increment when co-cultured with Δ*xac*2611 and a 10,000-fold increment when co-cultured with Δ8Δ2609Δ*xac*2611. In contrast, the viability ratio of the competition experiment between wild-type and Δ8Δ2609 *X. citri* strains was near unity, with neither strain outperforming the other, as previously shown (Oka et al. 2024). Competition experiments performed using *X. citri* cells in which the selective markers were inverted gave very similar results (**Figure 5**).

**Figure 5.**
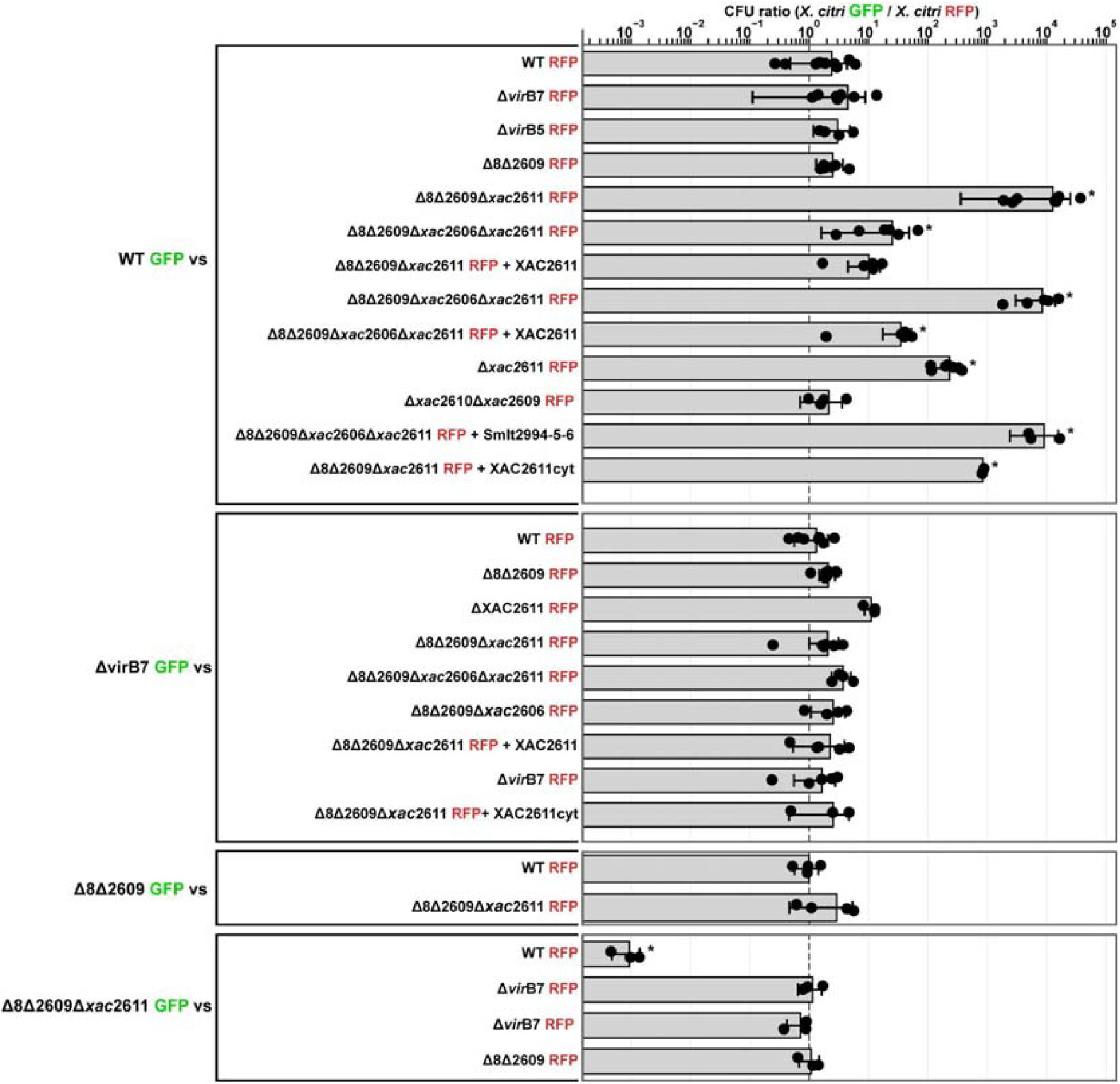
Quantitative analysis of X-T4SS-dependent fratricide in *X. citri* co-culture competition assays. For each competition assay, an equal number of cells of two *X. citri* strains carrying pBBR-2RFP plasmid, conferring kanamycin resistance, or pBBR-5GFP plasmid, conferring gentamicin resistance, were mixed and co-cultured at room temperature for two days on LB agar plates. Subsequently, colonies were resuspended, and the number of colony forming units (CFUs) for each strain was quantified using selective media. The bars in the figure depict the average CFU ratios, with individual dots representing the gentamicin-to-kanamycin ratio for each assay. Mean values and standard deviations (SD) were derived from 3 to 9 independent trials. Statistical comparisons were performed on log10-transformed CFU ratios using one-way ANOVA followed by Tukey HSD multiple-comparison testing. Asterisks indicate significant differences relative to the WT GFP versus WT RFP control condition, defined as Tukey-adjusted *p* < 0.05. The complete dataset, recalculated summary statistics, ANOVA results, Tukey HSD adjusted *p*-value matrix, significance matrix, and pairwise comparison table are provided in Supplementary Material S1.

We then tested the possible contribution of XAC2606, the second DUF4189 protein coded by *X. citri* X-T4SS gene cluster. The wild-type strain displayed a 25-fold viability increment in competition experiments with the Δ8Δ2609Δ*xac*2606 strain (**Figure 5**). Additionally, in competitions using the *X. citri* Δ8Δ2609Δ*xac*2606Δ*xac*2611 strain, the viability ratio of the wild-type strain was approximately 7,000, very similar to that observed when challenged against the Δ8Δ2609Δ*xac*2611 (see above). Moreover, when *X. citri* strains Δ8Δ2609Δ*xac*2606Δ*xac*2611 and Δ8Δ2609Δ*xac*2611, both transformed with a plasmid encoding XAC2611, were pitted against the wild-type, the competitive edge of the wild-type strain was markedly diminished (**Figure 5**). These results again confirm that the protection provided by XAC2606 against *trans*-intoxication is significantly less than that mediated by XAC2611 under the conditions tested.

A viability ratio close to unity was observed for competitions between the wild-type strain and the Δ*xac2610*Δ*xac2609* strain in which the glycoside hydrolase effector X-Tfe^XAC2609^ and its cognate immunity protein X-Tfi^XAC2610^, consistent with previous observations that the toxicity of the cocktail of X-Tfes secreted by the X-T4SS is resistant to the inactivation of individual effectors (Oka et al. 2024). Finally, competition assays using Δ8Δ2609Δ*xac*2606Δ*xac*2611 strains complemented with a cytoplasmic version of XAC2611 lacking its N-terminal signal peptide, failed to restore the competitiveness of these strains. The above results suggest that: (i) an intact XAC2611 is the primary factor preventing *trans*-intoxication in *X. citri* monocultures and (ii) XAC2611 must localize to the cell envelope to exert its protective function.

### The protective function of XAC2611 is species-specific

As shown in **Figure 1**), genes coding for XAC2611 homologs can be found within or adjacent to the loci coding for X-T4SSs in many other Lysobacteriales species. The *Stenotrophomonas maltophilia* K279 chromosome codes for three XAC2611 homologs (Smlt2994, Smlt2995 and Smlt2996) immediately downstream of the *virB6* gene (**Figure 1A**). We have previously shown that *X. citri* and *S. maltophilia* K279 cells in co-culture mutually kill each other in an X-T4SS-dependent manner (Bayer-Santos et al. 2019). This immediately suggests that XAC2611 protects against *trans*-intoxication mediated by the *X. citri* X-T4SS but not against the more distantly related *S. maltophilia* X-T4SS. To investigate this phenomenon further, we transformed the *X. citri* Δ8Δ2609Δ*xac2*606Δ*xac*2611 strain with a plasmid that co-expresses Smlt2994, Smlt2995 and Smlt2996. **Figure 5**) shows that the three *S. maltophilia* proteins were unable to protect against attack by the *X. citri* wild-type strain.

### Smlt2994 and Smlt2996 protect *X. citri* from attack by *S. maltophilia*

We then asked whether the heterologous expression of Smlt2994, Smlt2995 and Smlt2996 could protect *X. citri* cells against attack by *S. maltophilia* K279. **Figure 6A** shows colony co-culture competition assays between *S. maltophilia* wild-type or Δ*virD4* attacker strains and *X. citri* Δ8Δ2609Δ*xac*2606Δ*xac*2611 prey cells constitutively expressing GFP and carrying either an empty pBRA vector, the vector expressing XAC2611 (pBRA-XAC2611), one of the *S. maltophilia* XAC2611 homologs (pBRA-Smlt2994, pBRA-Smlt2995, pBRA-Smlt2996), or all three *S. maltophilia* genes in tandem (pBRA-Smlt299(4-5-6)). As expected, *S. maltophilia* out-competed *X. citri* Δ8Δ2609Δ*xac*2606Δ*xac2611* cells in an X-T4SS-dependent manner and this competitive advantage is maintained even when the *X. citri* Δ8Δ2609Δ*xac*2606Δ*xac2611* cells are complemented with the plasmid expressing XAC2611. However, *X-citri* cells regain their competitive advantage when expressing Smlt2994 or Smlt2996 individually or when expressing all three XAC2611 homologs (**Figure 6A**). Interestingly Smlt2995 did not confer protection against *S. maltophilia* attack. This is most likely due to the observed C-terminal truncation in this paralog (**Figure 1B**).

**Figure 6.**
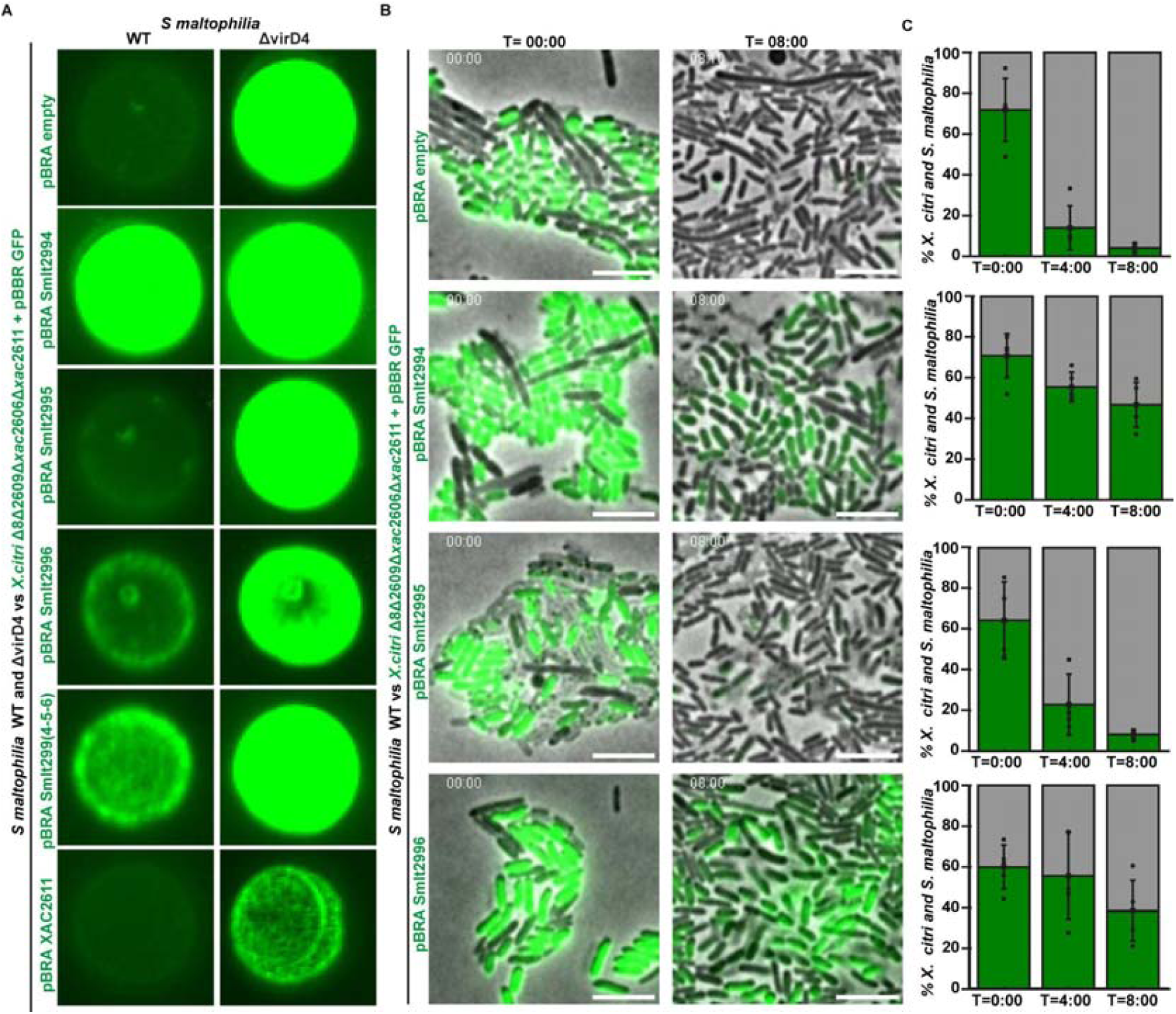
- Colony competition assays between *X. citri* and *S. maltophilia*. (A) Colony competition between *S. maltophilia* wild-type or Δ*vir*D4 attacker strains and *X. citri* Δ8Δ2609Δ*xac*2606Δ*xac2611* prey cells expressing constitutive GFP and carrying either an empty pBRA vector, pBRA-Smlt2994, pBRA-Smlt2995, pBRA-Smlt2996, the pBRA plasmid encoding the entire operon *smlt2994–smlt2995–smlt2996* [pBRA-Smlt299(4-5-6)], or pBRA-XAC2611. Mixed cultures were spotted on AB agarose medium, incubated for 7 days at room temperature, and fluorescence images were acquired using a Bio-Rad ChemiDoc MP imaging system. (B) Time-lapse fluorescence microscopy of *X. citri* versus *S. maltophilia* competition. *X. citri* Δ8Δ2609Δ*xac*2606Δ*xac2611 GFP (green cells)* carrying an empty pBRA vector, pBRA-Smlt2994, pBRA-Smlt2995, or pBRA-Smlt2996 are shown. Two frames from each movie are displayed, corresponding to the start (T = 00:00 h) and final (T = 08:00 h) time points. Additional examples of competition are shown in Supplementary Movies X. Strains and plasmid constructs used in each assay are indicated in the figure labels. Scale bars, 5 µm. (C) Quantitative analysis of the bacterial competition observed in **Supplementary Movies S14–S17**. Bars represent the mean percentage of visually identified *X. citri* GFP-positive cells and *S. maltophilia* cells at the onset of the experiment (T = 00:00 h), after 4 h (T = 04:00 h), and after 8 h (T = 08:00 h), scored across four to five representative microscopy fields (18 µm × 18 µm). Symbols denote values for each field, bars indicate the mean, and error bars represent the standard deviation. Across all conditions, a total of 1,128, 1,717, and 2,220 cells (*X. citri* plus *S. maltophilia*) were scored at T = 0, 4, and 8 h, respectively: for *X. citri* pBRA-Smlt2994 versus *S. maltophilia*, 313, 453, and 496 cells; for *X. citri* pBRA-empty versus *S. maltophilia*, 331, 570, and 721 cells; for *X. citri* pBRA-Smlt2995 versus *S. maltophilia*, 230, 265, and 451 cells; and for *X. citri* pBRA-Smlt2996 versus *S. maltophilia*, 254, 429, and 552 cells at T = 0, 4, and 8 h, respectively. At T = 00:00 h, the proportion of *X. citri* cells did not differ significantly among strains (one-way ANOVA, p = 0.51). By T = 04:00 h and T = 08:00 h, two clearly distinct groups emerged: a protected group, composed of *X. citri* Δ8Δ2609Δ*xac*2606Δ*xac2611* GFP cells expressing pBRA-Smlt2994 or pBRA-Smlt2996, which maintained ∼55% and ∼38–47% of the mixed population, respectively; and a susceptible group, composed of cells carrying the empty pBRA vector or pBRA-Smlt2995, which declined sharply to ∼14–23% and ∼4–8%, respectively (one-way ANOVA, p = 5.1 × 10[[ and p = 7.0 × 10[[; Tukey’s test, p = 0.0021–0.021 and p = 2.5 × 10[[–0.0017, at T = 04:00 h and T = 08:00 h, respectively). No significant differences were detected between strains within either group at any time point, indicating that Smlt2994 and Smlt2996 confer equivalent and robust protection, while Smlt2995 fails to protect *X. citri* from *S. maltophilia* X-T4SS-mediated killing. Scale bars (5 µm)

The results observed at a macroscopic scale (colonies) in **Figure 6A** were confirmed when observing the *X. citri* - *S. maltophilia* interactions at the cellular level using time-lapse fluorescence microscopy level over an 8 h period (**Figure 6B** and C and **Supplementary Movies S14–S17**). *X. citri* Δ8Δ2609Δ*xac*2606Δ*xac2611* expressing GFP and *S. maltophilia* cells were mixed at 1:1 proportions at the start of these experiments. During co-culture with *S. maltophilia*, *X. citri* cells carrying the empty vector were progressively eliminated, with extensive lysis by 4 h and near-complete elimination by 8 h. *X. citri* cells expressing Smlt2995 were likewise eliminated to a similar extent. In contrast, cells expressing Smlt2994 or Smlt2996 persisted throughout the experiment and remained abundant at 8 h. Thus, Smlt2994 and Smlt2996, but not Smlt2995, protected *X. citri* against *S. maltophilia*-mediated intoxication. Taken together, these results indicate that Smlt2994 and Smlt2996, but not Smlt2995, can functionally replace XAC2611 in the context of *S. maltophilia* attack.

### XAC2611 binds to VirB5

The results presented so far demonstrate that XAC2611 localizes throughout the cell envelope and prevents X-T4SS-mediated fratricide in a species-specific manner. We hypothesized that XAC2611 inhibits fratricide by directly interacting with an extracellular subunit of the attacking cell’s X-T4SS machinery. In characterized T4SSs, the major extracellular components include the major pilin VirB2 and the minor pilin VirB5 and the antenna domain of the VirB10 subunit of the outer membrane core complex (Schmidt-Eisenlohr et al. 1999; Fronzes et al. 2009; Lai and Kado 1998; Sgro et al. 2018). Both VirB2 and VirB5 have been hypothesized to function as adhesins, mediating initial contact with target or host cells (Yeo et al. 2003) and VirB5 has been shown to localize at the tip of the T4SS pilus in specific systems (Aly and Baron 2007). Finally, in the structure and functional model of the pKM101 conjugative T4SS described by (Macé et al. 2022; Macé and Waksman 2024), VirB5 is proposed to emerge from the outer membrane pore of the T4SS at the distal end of the growing pilus, suggesting a role in interactions with target cell. Thus VirB5 is a strong candidate as a possible interaction partner with XAC2611.

To test whether XAC2611 interacts directly with VirB5, we performed pull-down assays using lysates of *E. coli* cells expressing residues 51–275 of *X. citri* VirB5 fused to a C-terminal S-tag and/or residues 55–164 of XAC2611 fused to an N-terminal His-tag. Both constructs lacked their N-terminal signal peptides and predicted intrinsically disordered N-terminal regions. **Figure 7**) shows that His-XAC2611_(55–164)_ bound effectively to a Ni²[-NTA column, whereas VirB5_(51–275)_-S-tag alone did not. However, VirB5_(51–275)_-S-tag was retained on the column in the presence of His-XAC2611_(55–164)_ (**Figure 7**)). These results confirm a direct interaction between XAC2611 and VirB5.

**Figure 7.**
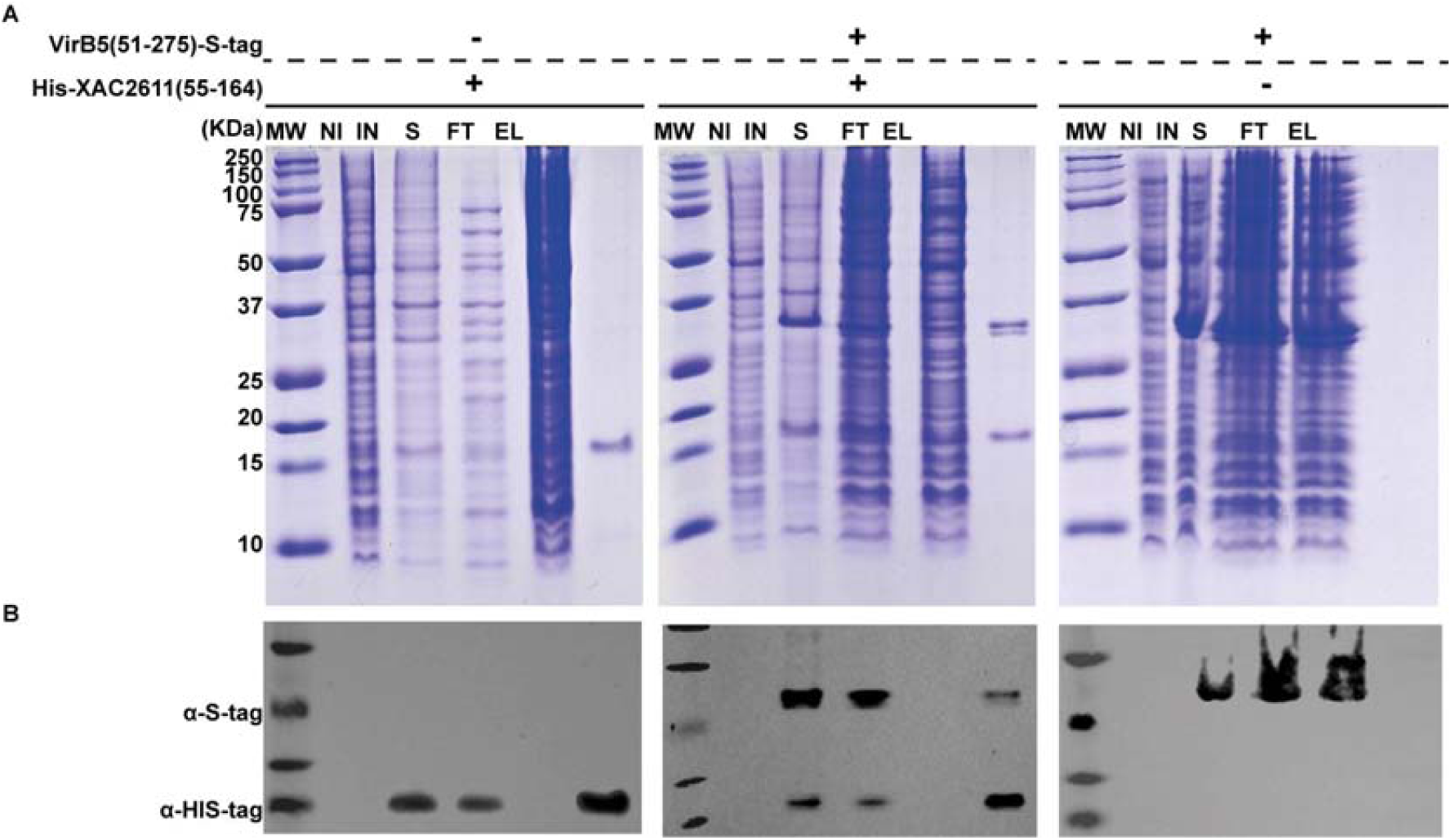
XAC2611 interacts with VirB5. **(A)** Coomassie-stained 16% Tricine-SDS-PAGE showcasing Co^2+^-NTA affinity chromatography pulldown assay between His-XAC2611 and VirB5-S tag. *E. coli* BL21De3 harbored pRSF duet encoding His-XAC2611(55-164) (left column), both His-XAC2611(55-164) and VirB5(51-275)-S tag (middle column), or VirB5(51-275)-S tag alone (right column). Total cell lysates of protein expression before (NI) and after induction (IN) of recombinant proteins, soluble protein fractions (S) prior to loading into the Co^2+^-NTA column, flow-through (FT) of unbound proteins washed with buffer A (20 mM Tris-HCl pH 7.3, 200 mM NaCl, 20 mM imidazole, 20% glycerol) and elution fraction (EL) using an imidazole gradient (20 mM - 500 mM). Molecular weight standards (MW). **(B)** Western blot analysis of the fractions presented in part (A) using primary anti-polyhistidine antibody (α-HIS-tag) or anti-S-tag antibody. Detected fractions correspond to the lanes displayed in part (A) of the figure.

## Discussion

We have shown that the DUF4189 encoding-protein XAC2611 is the key factor preventing *trans*-intoxication via the X-T4SS among sibling *X. citri* cells. XAC2611 provides kin-specific protection in *X. citri* by selectively inhibiting the transfer of antibacterial effectors among genetically identical sibling cells. This specificity is highlighted by the observations that the homologous proteins from *S. maltophilia* protect *X. citri* ΔXAC2611 mutants against attack by *S. maltophilia* cells (**Figure 6**)) but not against attack by wild-type *X. citri* cells (**Figure 5**)). Thus, XAC2611 acts as a molecular safeguard, inhibiting the delivery of toxic effectors from neighboring kin and preserving cellular integrity within clonal populations. We therefore propose to name XAC2611 and its orthologs as *trans*-intoxication protein factors (Tpfs; for example Tpf^XAC2611^ in *X. citri* and Tpf^Smlt2994^ and Tpf^Smlt2996^ in *S. maltophilia*).

We demonstrated that Tpf^XAC2611^ can specifically interact with VirB5 within the complex mixture of an *E. coli* cell lysate. Structural and biochemical studies of VirB5 from diverse organisms have established that it is located at the distal tip of the T4SS pilus, functioning as an adhesin that facilitates contact with recipient cells (Macé et al. 2022; Schmidt-Eisenlohr et al. 1999; Lai and Kado 1998; Yeo et al. 2003; Aly and Baron 2007). Based on the structural information on the T4SS (Macé et al. 2022) and our data, we propose a model (**Figure 8**)) whereinTpf^XAC2611^, localized in the periplasm of recipient cells, directly interacts with the VirB5 component at the tip of the T4SS pilus from attacking sibling cells. This interaction may block the formation of a mature and stable junction between donor and recipient cells, thus inhibiting the translocation of antibacterial effectors. Unlike paradigmatic models which posit exclusively on the presence of cognate intracellular immunity proteins to neutralize toxic effectors translocated from kin cells (Jurėnas and Journet 2021; Russell et al. 2011), the model presented in **Figure 8**) proposes that obstruction of the secretion machinery at the point of contact between donor and recipient cells prevents effector delivery altogether.

**Figure 8.**
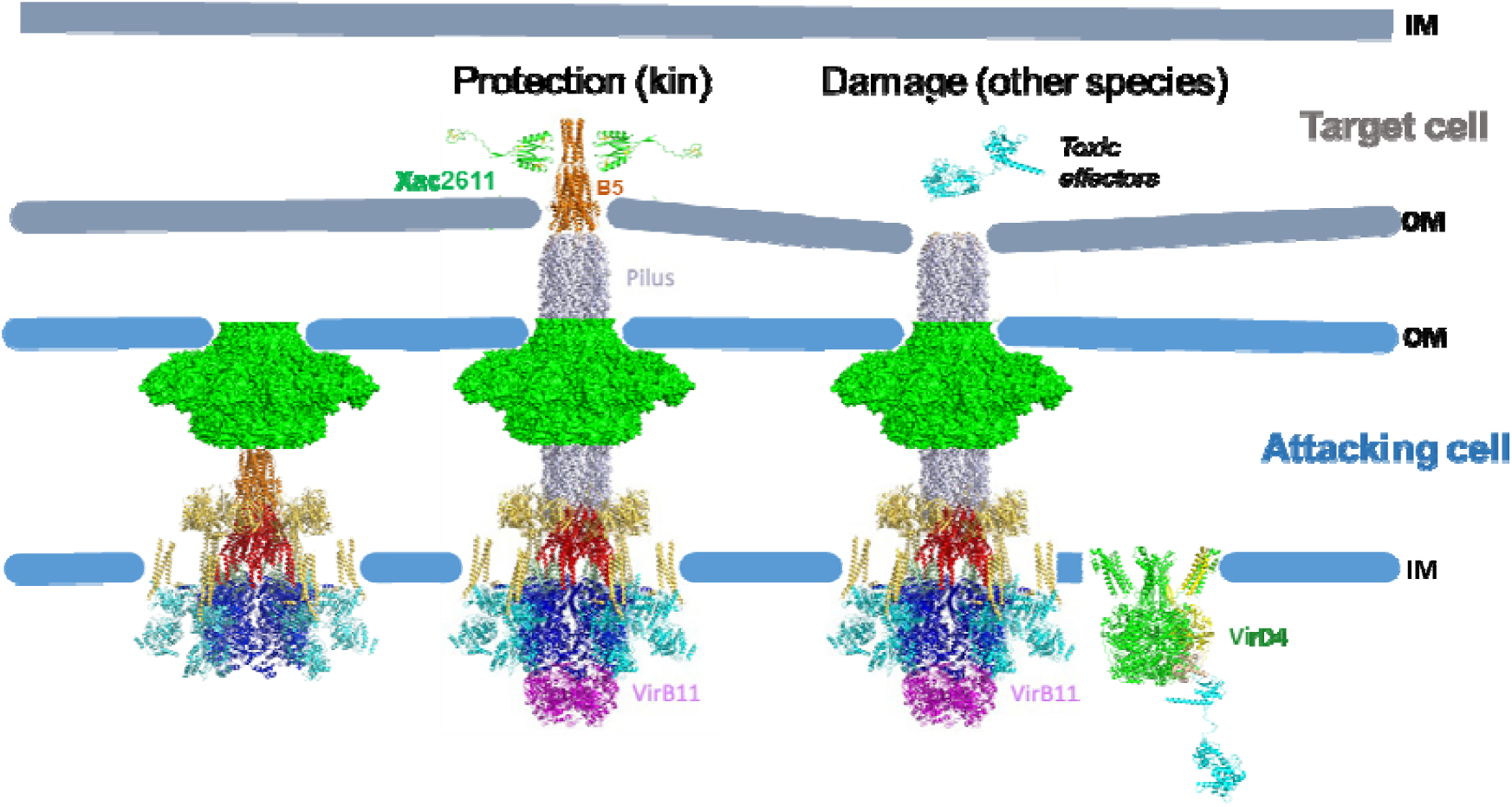
Model for kin-specific protection mediated by the X-T4SS in *X. citri.* Schematic representation of effector delivery outcomes in the presence (left, “Protection”) or absence (right, “Damage”) of the periplasmic exclusion protein XAC2611 in the recipient cell. The structural model of the Type IV Secretion System (T4SS) includes the core complex (VirB3–VirB10), the coupling protein VirD4, and the minor pilin VirB5 located at the tip of the pilus. In the “Protection” scenario, XAC2611 binds to VirB5, preventing engagement with the target cell and thereby blocking effector translocation. In the “Damage” scenario, the absence of XAC2611 allows docking of the pilus onto the recipient cell envelope, enabling translocation of toxic effectors into the periplasm via the VirD4-coupled secretion machinery (Oka et al. 2022). IM, inner membrane; OM, outer membrane. The T4SS structural model is based on the conjugative T4SS architecture (Macé et al. 2022), and XAC2611 was modeled using AlphaFold3 (Abramson et al. 2024).

One outstanding question is what molecular mechanism underlies the action of VirB5 in the target cell and how does XAC2611 and its homologs interfere with this mechanism? In the Mace et al (2022) and Mace and Waksman (2024) cryoEM structure of the conjugative pKM101 T4SS, the VirB5 pentamer is docked on top of the VirB6 platform in the absence of the VirB2 pilus in what could be interpreted as a pre-activated state within the periplasm. However, VirB5 may undergo significant structural changes upon pilus extension and interaction with the target cell envelope. It is in this post-activated state that the Tpf^XAC2611^ -VirB5 interaction may become physiologically relevant.

The Tpf^XAC2611^ protein is 164 residues in length. It belongs to a large homologous superfamily of proteins (DUF4189) predicted to adopt a conserved globular topology stabilized by three highly conserved disulfide bonds, consistent with the predicted and observed localization in the oxidative environment of the periplasm. In many cases the genes of Tpf^XAC2611^ orthologs are unannotated and may occur in multiple copies and almost all XAC2611 orthologs associated with X-T4SSs carry a signal sequence for transport to the cell envelope (**Figure 1**)). This signal sequence is predicted to be cleaved by the signal peptidase (Teufel et al. 2022) and is followed by a proline-rich region containing another pair of conserved cysteine residues predicted to form an intrinsically disordered N-terminal tail that precedes the globular domain (**Figure 1**)).

Our proposed mechanism of a cell envelope protein blocking the transfer of X-T4SS substrates is analogous to the phenomenon of “self-incompatibility” in plasmid conjugation that has been known since the 1950s (Lederberg et al. 1952; Kingsman and Willetts 1978; Arutyunov and Frost 2013). Proteins involved in surface and entry exclusion phenomena have been proposed to act via several mechanisms in different conjugative systems, such as interactions between TraS and TraG (Audette et al. 2007; Anthony et al. 1999) and Eex and TraG in F-like plasmid and ICE systems (Marrero and Waldor 2005), respectively. The Tpf^XAC2611^-VirB5 interaction described here is the first example that we know of demonstrating a direct physical interaction between a cell envelope-associated exclusion protein and its molecular target in the T4SS. It would be interesting to test if similar interactions between exclusion proteins and VirB5 homologs occur in other T4SSs. Interestingly the sequences of Tpf and VirB5 proteins associated with X-T4SS show very high sequence variability (see Tpf alignments in **Figure 1**) and VirB5 alignments in (Sgro et al. 2019)) consistent with their important roles in distinguishing kin (friend) from non-kin (foe).

Further exploration of bacterial genomes reveals a broad distribution of DUF4189 (InterPro: PR025240) proteins across diverse taxonomic groups, including Alphaproteobacteria, Gammaproteobacteria, Betaproteobacteria, Actinomycetes, and Cyanophyceae. Species within the Gammaproteobacteria and Betaproteobacteria classes, where DUF4189 is commonly found, are often but not solely associated with the presence of a chromosomally encoded X-T4SS (Souza et al. 2015; Oka et al. 2022). In Actinomycetes, the DUF4189 domain is found in the Mycobacterial effector Rv1813c, secreted by *Mycobacterium tuberculosis* and acts in the mitochondria where it modulates host cell metabolism (Martin et al. 2023). Structural analyses of Rv1813c reveal a conserved core fold composed of two tandem βββαβ motifs stabilized by three disulfide bonds in a manner equivalent to that predicted for Tpf^XAC2611^ and its homologs. Despite high sequence variability in the DUF4189 superfamily (**Figure 1**)), the structural integrity of the fold is evolutionarily maintained, and possibly strengthened by the predicted highly conserved dissulphide bonds. Such conservation likely supports a stable scaffold capable of functional diversification across taxa.

---

**Supplementary Figure 1.**
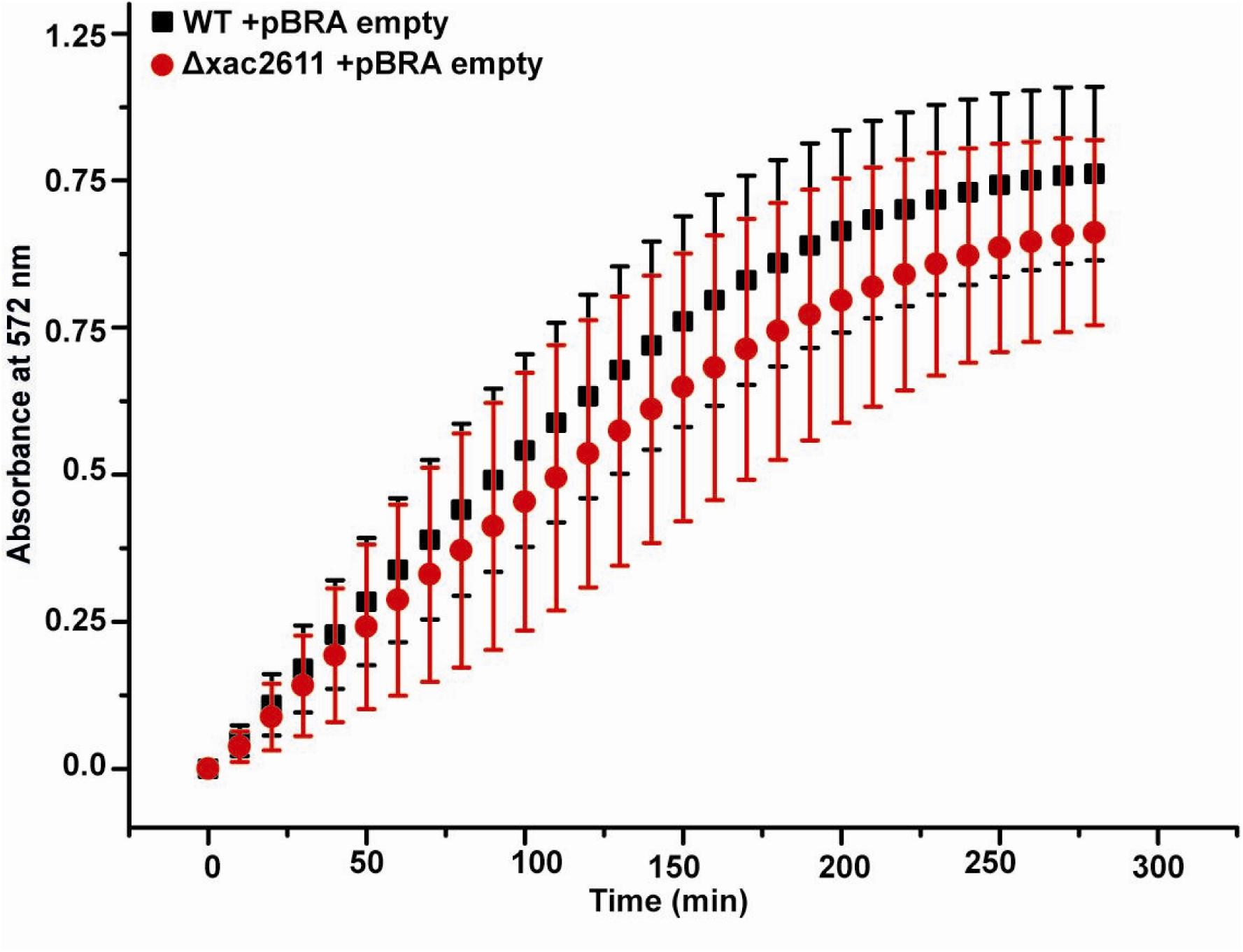
CPRG Interbacterial competition assay between *X. citri* and *E. coli* MG1655 expressing β-galactosidase, monitored by CPRG assay in 96-well plates filled with agarose media using a microplate reader. *X. citr*i strains include wild type (WT), mutant and Δ*xac2611* carrying empty plasmid pBRA (pBRA empty). Mean and SD of 5 independent assay.

**Supplementary Figure 2.**
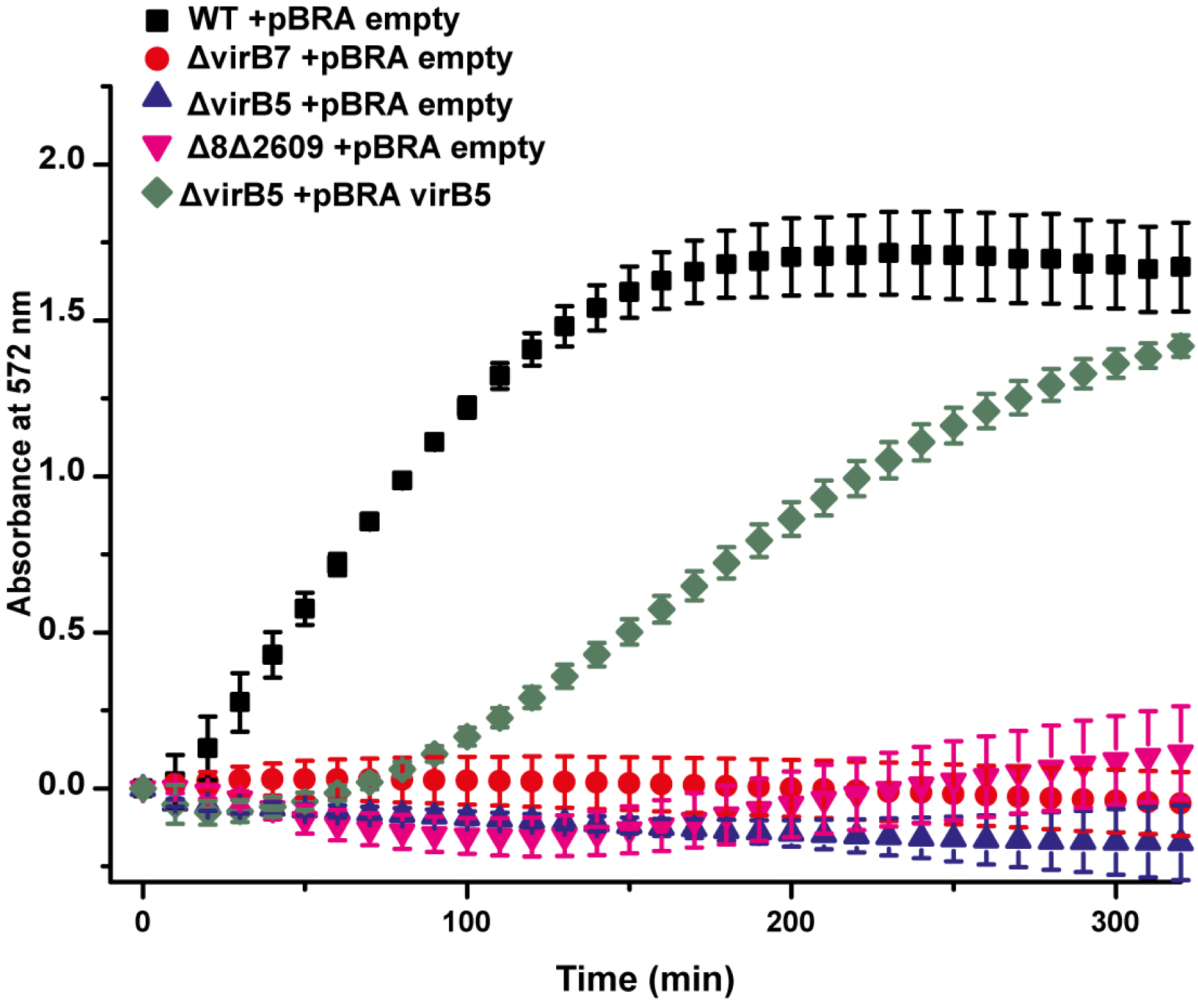
CPRG Interbacterial competition assay between *X. citri* and *E. coli* MG1655 expressing β-galactosidase, monitored by CPRG assay in 96-well plates filled with agarose media using a microplate reader. *X. citr*i strains include wild type (WT), mutants Δ*virB7* and Δ8Δ2609-GFP (Δ8Δ2609) carrying empty plasmid pBRA (pBRA empty), and Δ*virB5* strain carrying pBRA empty, pBRA encoding virB5 (pBRA *virB5*).

## Material and methods

### Bacterial strains and cloning

Oligonucleotides, plasmids, and bacterial strains used in this work are listed in Tables S1, S2, and S3, respectively. *Xanthomonas citri* strain 306 (hereafter *X. citri*) was typically grown as previously described (Oka et al., 2024). Briefly, the antibiotics employed were ampicillin (100 μg/mL), kanamycin (50 μg/mL), gentamicin (20 μg/mL), and spectinomycin (100 μg/mL). Cell cultures were generally grown on LB medium solidified with 1.5% (w/v) agar and supplemented with the appropriate antibiotics at 30 °C for 24–48 h. Isolated colonies were picked and grown overnight in 2×TY medium at 30 °C with shaking at 200 rpm. The optical density at 600 nm (OD[[[) of the cultures was then adjusted to 0.05 in 2–5 mL of fresh 2×TY medium and cells were grown at 30 °C, 200 rpm for an additional 14–18 h before the assays. *Stenotrophomonas maltophilia* wild-type and ΔvirD4 strains were typically cultured in 2xTY medium at 37 °C, 200 rpm as overnight pre-inocula, diluted 50-fold, and grown for 4 h before the assays, as described in Bayer-Santos et al. (2019).

### Cloning and Generation of X. citri Gene-Knockout Strains

The genomic DNA sequence of X. citri strain 306 (da Silva et al. 2002) and *S*. *maltophilia* K279a was employed as a template for PCR amplification using the oligonucleotides listed in Table S1. The resulting amplicons were incorporated into plasmids pBRA (M. Marroquin, unpublished) and pRSF duet to generate pNPTS-derived suicide vectors (Table S2) using restriction enzymes following the reaction of ligases or Gibson Assembly (NEB) reactions. Deletion of X. citri genes was performed as described in (Oka et al. 2022). Briefly, pNPTS-derived transformant *X. citri* cells selected in LB agar supplemented in kanamycin and susceptible growing in LB-agar supplemented with 5% sucrose were submitted to grown overnight in 2xTY media without antibiotics and plated in LB agar plates supplemented with 5 % sucrose. Then, clones susceptible to growing in LB plates supplemented with kanamycin and viable in sucrose were selected for genomic PCR to check the deletion of the genes of interest.

To obtain plasmids pRSF DuetMCS1-XAC2611(55-164) and pRSF DuetMCS1-XAC2611(55-164)-MCS2-VirB5(51-275), a synthetic gene encoding HIS XAC2611(55-164) into pET28a-HIS XAC2611(55-164) was transferred into pRSF derived vectors using the NcoI-BamHI restriction sites. Cloning sequences and plasmid sequences were verified by DNA sequencing.

### Time-lapse fluorescence and Immunofluorescence microscopy

All time-lapse fluorescence and immunofluorescence microscopy procedures were conducted at room temperature, except for the cell growth phase in liquid 2×TY broth, which was carried out at 30 °C to obtain cell cultures. Time-lapse fluorescence microscopy assays followed established protocols (Oka et al., 2022, 2024). Briefly, overnight cultures of *X. citri* and *S. maltophilia* were pelleted by centrifugation in 1.5 mL microtubes (5,000 rpm, 5 min), washed twice with 2×TY, and normalized to an OD[[[ of 2.0. Then, 1 µL of each *X. citri* strain, or 1:1 mixtures used for competition assays (*X. citri* vs. *X. citri*), was deposited onto a thin LB–agarose pad and covered with a coverslip. Mixtures of *X. citri* and *S. maltophilia* (1:1 ratio) were spotted onto AB–agarose pads and similarly covered with coverslips. Time-lapse microscopy of isolated *X. citri* cells and bacterial competition was performed using a Nikon Eclipse Ti microscope equipped with filters for GFP (GFP-3035B-000-ZERO, Semrock) and RFP (TxRed-4040B, Semrock), along with a Nikon Plan Apo 100× objective. Images were acquired at 10 min intervals and processed using NIS-Elements software (version 3.07; Nikon). Competition assays involving *X. citri* vs. *S. maltophilia* were additionally observed with a Leica DMi8 microscope equipped with a GFP fluorescence filter (excitation 470 ± 40 nm, dichroic mirror 500 nm, emission 525 ± 50 nm; Leica). Time-lapse movies and images were processed and analyzed with Fiji (Schindelin et al., 2012), and drift was corrected using the StackReg plugin (Thévenaz, Ruttimann, and Unser, 1998). Statistical analyses were performed using OriginPro 2020 (OriginLab).

*X. citri* Δ*xac*2611 strain, transformed with plasmid pBRA-XAC 2611-FLAG, was cultivated for 12 hours in 2 mL of 2xTY medium at 30°C with shaking at 200 rpm. Subsequently, 1 mL of the culture was collected in a microtube, and the cells were centrifuged at 5000 rpm for 5 minutes. Then, cells were washed by discarding the supernatant and resuspending in 1 mL of PBS buffer (100 mM phosphate buffer pH 7.4, 300 mM NaCl). This wash step (pelleting, discarding the supernatant, and resuspending PBS) was repeated twice. Next, the cells were centrifuged (5000 rpm, for 5 minutes) and then resuspended in 1 mL of PBS buffer supplemented with 4% formaldehyde for 30 minutes. Next, cells were centrifuged (5000 rpm for 5 minutes) and washed with PBS. Then, 500 μL aliquots of the cells were distributed into two new microtubes. Cels were centrifugated (5000 rpm for 5 minutes), the supernatant was discarded, and each tube was resuspended in 70% ethanol or distilled water for 30 minutes. Subsequently, the cells were washed twice with PBS and resuspended for a final volume of 500 uL. Next, two aliquots of 250 μL derived from the previous treatment were distributed into four new microtubes, cells were centrifuged (5000 g, 5 minutes, room temperature), and the supernatant was discarded. The cultures were then resuspended in PBS supplemented with 1% BSA and lysozyme (1 µM) or in PBS supplemented with 1% BSA for 30 minutes at room temperature, followed by two washes with PBS supplemented with 1% BSA. Following the washing steps, the samples were incubated with primary antibodies for 30 minutes (1:1000) anti-FLAG (ANTI-FLAG® Merck, F7425), washed three times (PBS supplemented with 1% BSA and 0.1% Tween 20), and then with secondary antibodies (1:1000) (Alexa Fluor® 488) (ab150077) following two wash steps with PBS supplemented with BSA 1% and 0.1% Tween 20. Subsequently, cells were spotted in LB-agarose pads and observed with a Leica DMi-8 microscope equipped with a GFP fluorescence filter (excitation 470 +/-40, dichroic mirror 500 nm, emission 525+/-50, Leica). *X. citri*::XAC2611-GFP and *X. citri:*:XAC2606-GFP cell cultures were acquired using the same microscope and filters (Leica DMi-8 microscope equipped with a GFP fluorescence filter). Images were acquired using Z-stacking (10-30 steps of 50 nm). Blind deconvolutions for fluorescence channels were performed with Leica LAS X software. Subsequent analyses were performed using Fiji 2.

### Protein Purification Pull-Down Assays Western Blot

Cell colonies of *E. coli* Bl21De3 on LB agar plates transformed with plasmids pRSFDuet derivatives (Table SX) were typically inoculated in 5 mL of 2xTY broth medium and grown overnight (37 C, 200 rpm). Cell cultures were then diluted in 500 mL of 2xTY and allowed to grow at 37 °C until D.O600nm=0.7. Next, induction was followed by adding IPTG (0.5 mM) for 16 hours at 18 °C, 200 rpm. Then, the cells were pelleted by centrifugation, resuspended in lysis buffer (25 mM Tris-HCl (pH 7.2), 200 mM NaCl, 20% (v/v) glycerol), and lysed with a French press at 4 °C. The soluble fractions were recovered after centrifugation for 45 min at 25000 g. Affinity chromatography was performed using an FPLC AKTA system (Cytiva) with HisTrap HP 5 mL (GE) charged with Co^2+^ equilibrated with 25 mM Tris-HCl buffer (pH 7.2), 200 mM NaCl, 20 mM imidazole, and 20% (v/v) glycerol. Typically, sample loading, washing (75 mL), and elution (20 to 500 mM imidazole gradient, along 75 mL) were performed at 5 ml/min. Protein fractions co-eluted from the pull-down assay of VirB5(51-275)-S-tag with His-XAC2611(55-164) were resolved on Tricine (ref) SDS-PAGE 15% gel and transferred to a nitrocellulose membrane using a semi-dry blot system at 60 mA for 1 hour. Following the transfer, the membrane was blocked with TBS (25 mM Tris-HC, 100 mM NaCL|) buffer supplemented with 10% non-fat milk, agitated at 50 rpm for 16 hours, and then probed with anti-S-Tag rabbit antibody (ABCAM 183674) (1:1000 dilution) for two hours at room temperature at 50 rpm. Subsequently, the membrane was washed five times with TBS buffer supplemented with 10% non-fat milk and 0.1% tween 20, each wash lasting 5 minutes at 50 rpm, followed by incubation with IRDye 800CW goat anti-rabbit IgG (LI-COR Biosciences) (1:20000) dilution for 30 minutes at room temperature 50 rpm. Then, the membrane was washed with TBS, and Images were collected using the ChemiDoc System MP (BioRad). For detection of His-XAC11 Anti-polyhistidine-alkaline phosphatase conjugated (Sigma A5588) (1:20000 dilution, two hours, 50 rpm), was incubated for one hour with the nitrocellulose membranes, washed and revealed with 22.5 ul de BCIP (50 mg/ml) e 30 ul de NBT (75 mg/m 70 % dimethylformamide) in buffer (100 mM Tris-HCl, pH 9.5, 100 mM M NaCl, 5 mM MgCl), images were acquired using the ChemiDoc System MP (BioRad).

### X. citri vs X. citri co-culture competition assays

The experimental procedures followed the methodology outlined by (Oka et al. 2024). In brief, for the bacterial competition assay based on viability, X. citri strains carrying either plasmids pBBR-5GFP or pBBR-2RFP were cultured at 30 °C for 12 h in 2xTY media supplemented with gentamicin (20 µg/mL) or kanamycin (50 µg/mL), respectively. Following three washes with fresh 2xTY medium, cultures of *X. citri* at OD600nm of 2.0 were mixed in a 1:1 ratio (v:v). Five microliters of this mixture were then spotted onto 1% LB-agar plates containing ampicillin (100 µg/mL) and incubated for 2 days (40-50 hours) at room temperature (22 °C-25 °C). Subsequently, colonies were collected by resuspending them in 1 mL of 2xTY medium, and the viability of each colony was determined via serial dilution on selective media with kanamycin or gentamicin-supplemented LB-agar plates.

### *X. citri* vs *E. coli* competition assays

The bacterial competition assays based on Chlorophenol-red β-D-galactopyranoside (CPRG) were conducted using the same procedure described in (Oka et al. 2022). In brief, X. citri cultures with an OD600nm of 2.0 were mixed in a 1:1 (v:v) ratio with *E. coli* MG1655 cultures at OD600nm of 11. These mixtures (5 μL) were then applied to 1.5% agarose supplemented with 40 µg/mL CPRG (Sigma-Aldrich) in 96-well plates. Absorbance at 572 nm was recorded every 10[minutes using a plate reader (SpectraMax Paradigm, Molecular Devices).

### *X. citri vs. S. maltophilia* co-culture competition assays

Assays were performed as previously described in Bayer-Santos et al. (2019). Briefly, *S. maltophilia* cells from overnight cultures were washed three times with 2×TY medium and normalized to an OD of 0.05, then mixed at a 1:1 ratio with a washed culture of *X. citri* adjusted to an OD of 2.0. Five microliters of the mixed cultures were spotted onto AB medium (0.2% (NH) SO, 0.6% Na HPO, 0.3% KH PO, 0.3% NaCl, 0.1 mM CaCl, 1 mM MgCl, 3 μM FeCl) supplemented with 0.2% sucrose, 0.2% casamino acids, 25 μg/mL thiamine, and 0.5% arabinose, and solidified with 1% agarose. The colonies were monitored for seven days at room temperature (22 °C).

## Supporting Information

**S1 Table.**
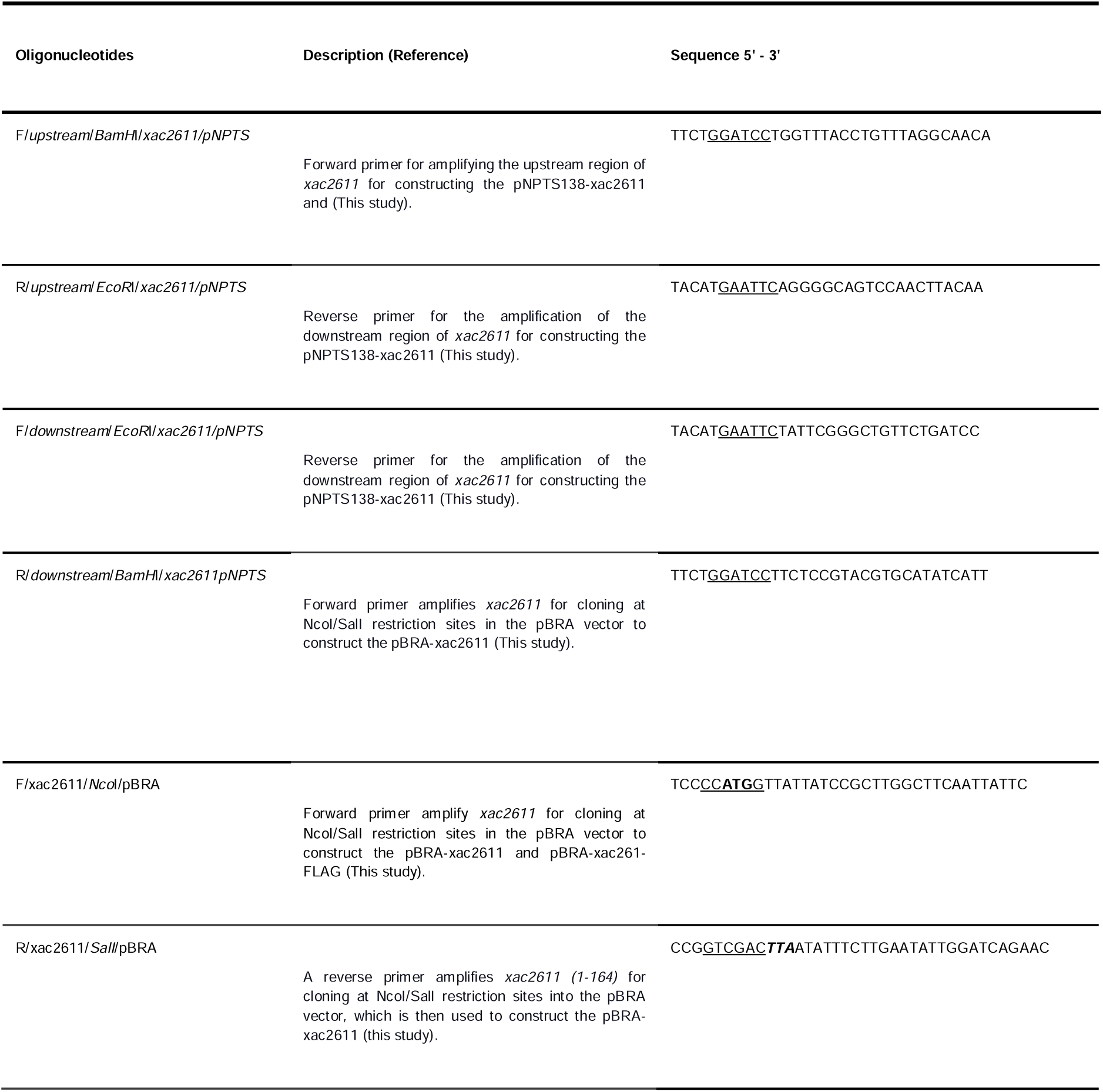
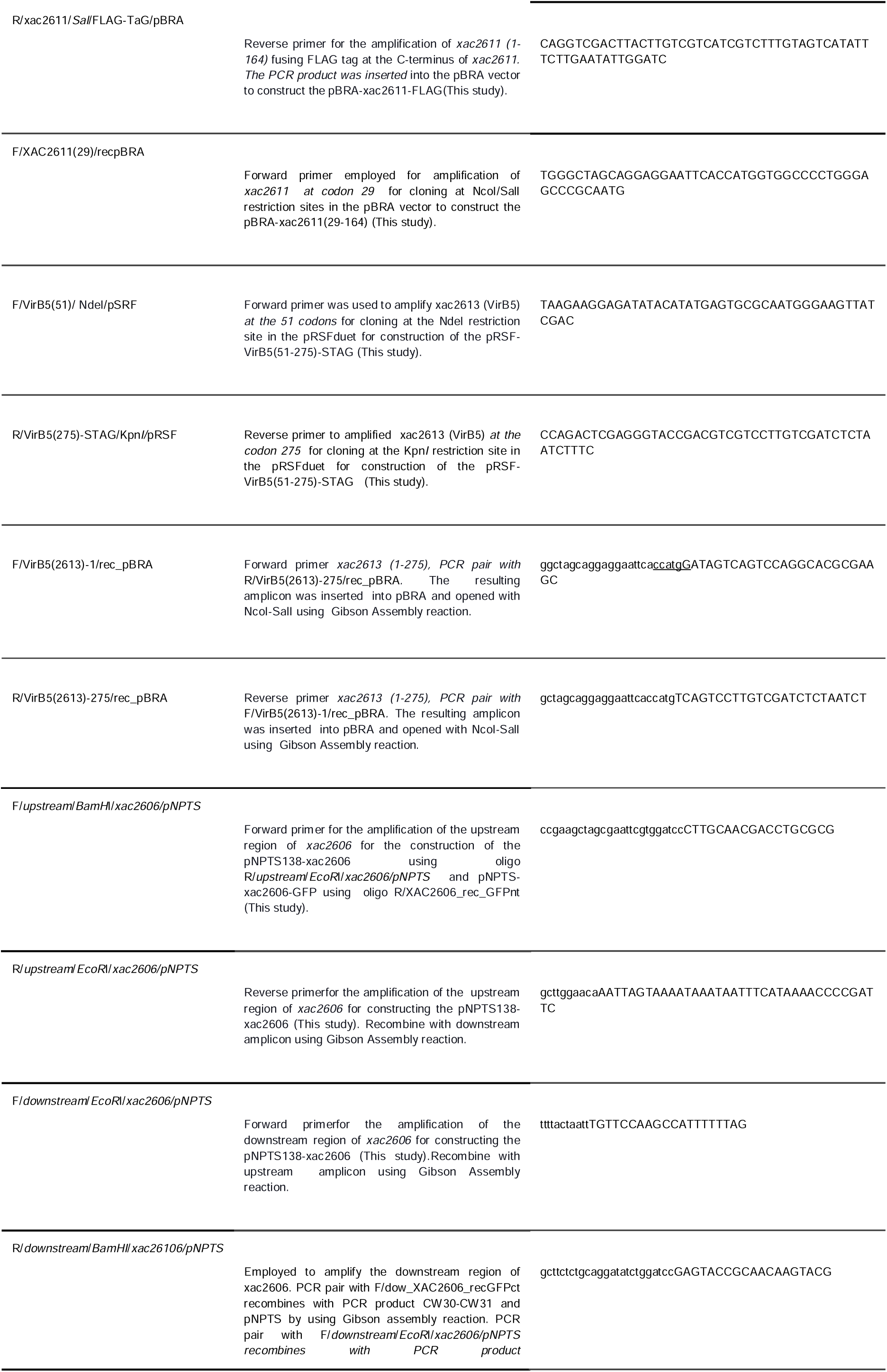
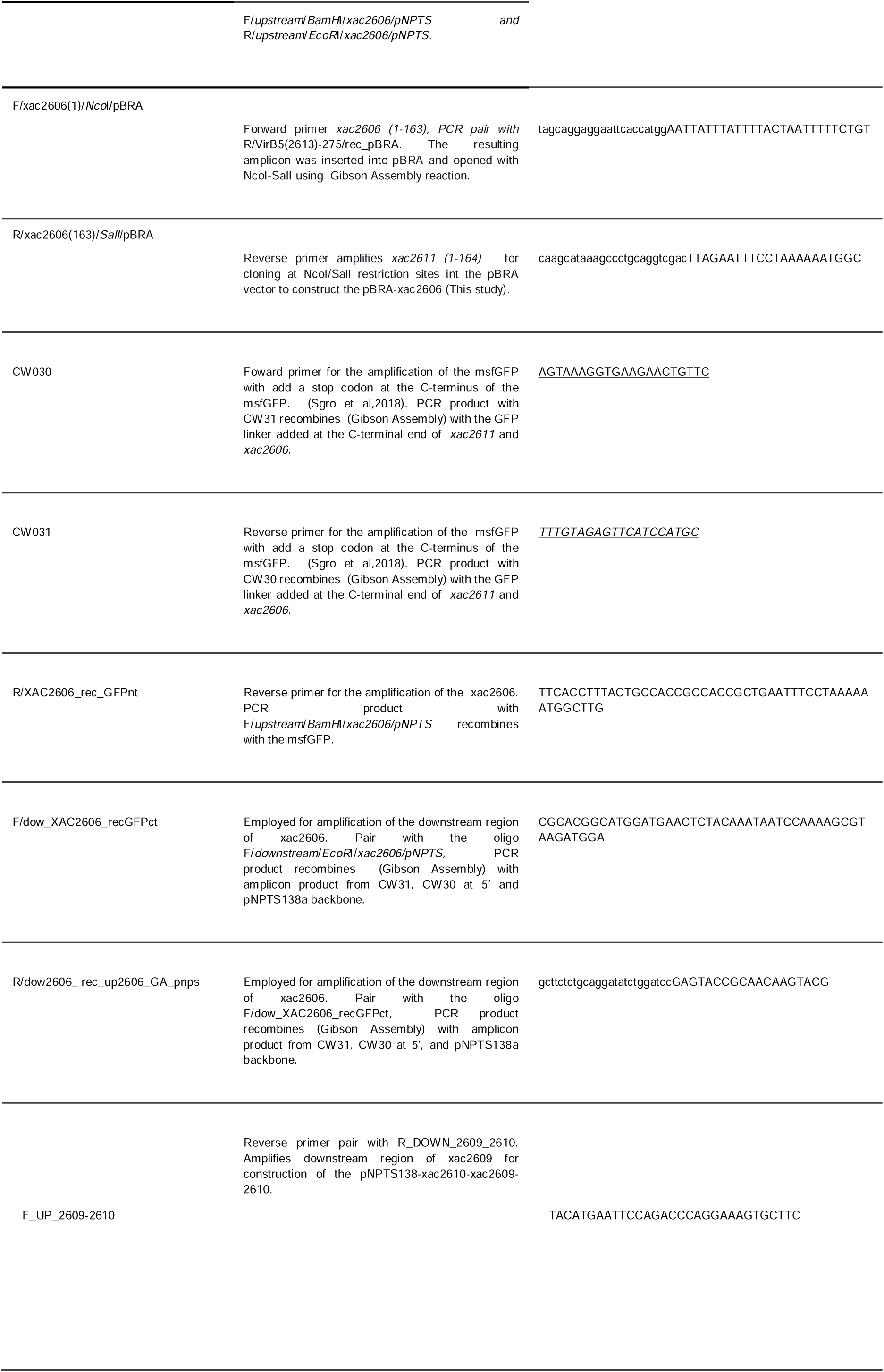
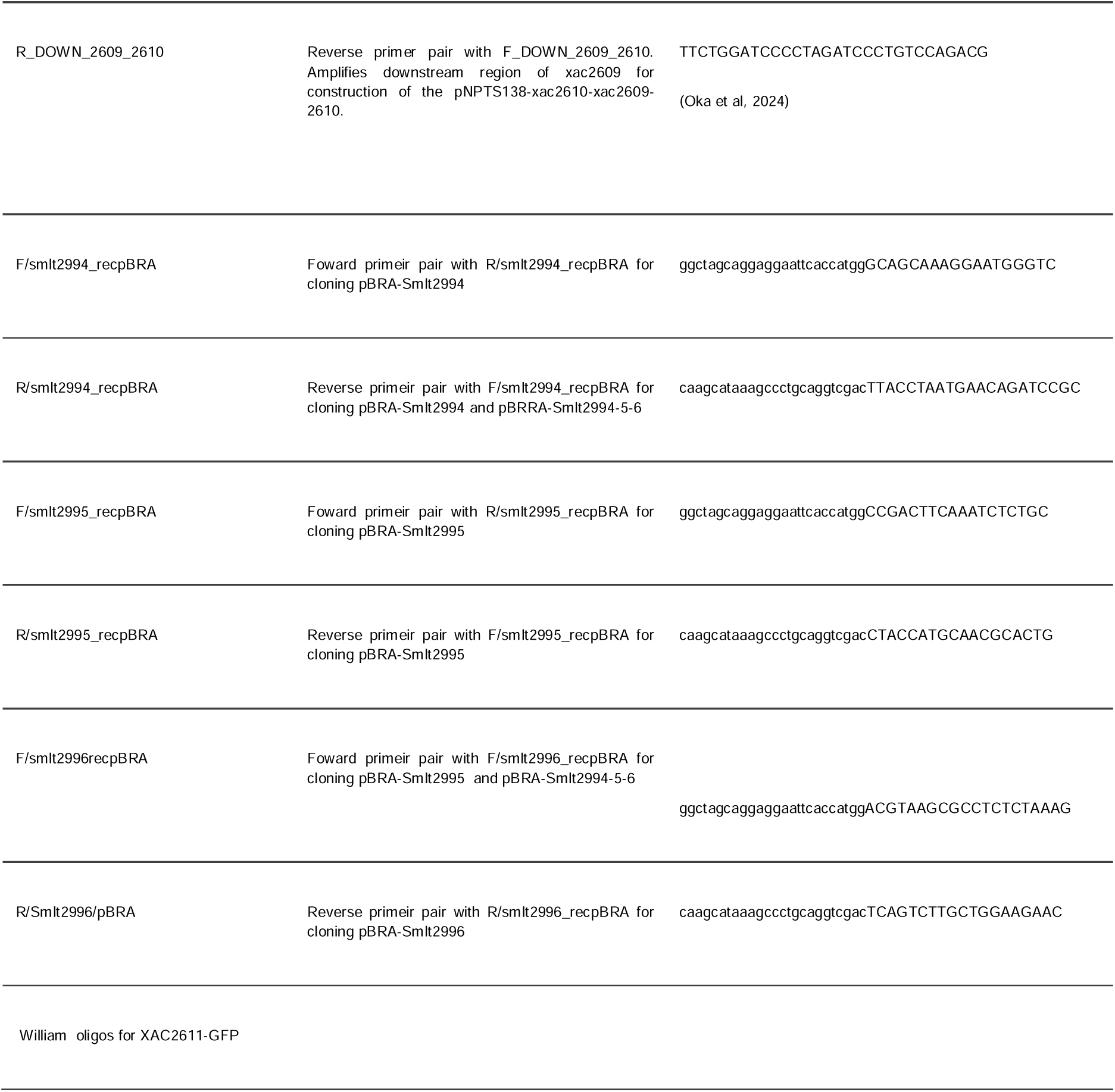
Oligonucleotides used in this study.

**Table S2.** Plasmids used in this study.

| plasmid | Properties | references |
| --- | --- | --- |
| pBRA | pBAD24 derivative vector, which has an arabinose-inducible promoter that is constitutively active plasmid in *X. citri*.Spc<sup>R</sup>. | M.Marroquin, unpublished |
| pBBR-5GFP | Broad host range plasmid cloning vector pBBR1MCS-5 expressing GFP; Gm | (Dunger et al. 2014) |
| pBBR-2RFP | Broad host range plasmid cloning vector pBBR1MCS-5 expressing RFP; Gmr | (Oka et al. 2022) |
| pNPTS138 | Suicide vector for generation of gene knockouts (sacB), Km<sup>r</sup> | M.R.Alley unpublished |
| pRSF-DUET | Expression vector (addgene) under control of lac operon, expression induced in E.coli BL21De3 strain by adding IPTG in the cell culture at the end of the exponential phase. This vector presents two multiple cloning sites. Km<sup>r</sup> |  |
| pET28a-HIS-XAC2611(55-164) | Synthetic construct (Genscript) codon optimized for expressing HIS6-XAC2611 in E.coli, Km<sup>r</sup>. Fragment encoding XAC2611(55-164) was excised with NcoI-BamHI and inserted into pRSF-DUET and pRSF-MCS2-VirB5-STAG5(1-275). | This work |
| pRSF-MCS1HIS6-XAC2611(55-164)-MCS2-VirB5-STAG(51-275) | pRSF-DUET coding for XAC2611(55-164) fused with a 6His( polyhistidine tag) at the N-terminus at the NcoI-BamHI sites in MCS and XAC2613 (51-275) fused with an S-tag at the C-terminus at the NdeI-KpnI sites of the MCS2. Km<sup>r</sup> | This work |
| pRSF-MCS1HIS6-XAC2611(55-164) | pRSF-DUET coding for XAC2611(55-164) fused with a 6His( polyhistidine tag) at the N-terminus at the NcoI-BamHI sites in MCS1. Km<sup>r</sup>. | This work |
| pRSF-MCS2-VirB5-STAG5(51-275) | pRSF-DUET coding for XAC2613 (51-275) fused with an S-tag at the C-terminus. Inserted at the NdeI-KpnI sites of the MCS2.Km<sup>r</sup>. | This work |
| pBRAXAC2611(1-164) | pBRA vector for the expression of XAC2611(1-164) ,xac2611 orf was inserted at the NcoI-SalI cleavage sites.Spc<sup>R</sup>. | This work |
| pBRAXAC2611(1-164)-FLAG | pBRA vector for expressing XAC2611(1-164) with a C-terminus FLAG sequence. | This work |
| pBRAXAC2611(29-164) | pBRA vector for the expression of XAC2611(29-164) (without a signal peptide and lipobox) XAC2611(29-164)f was inserted at the NcoI-SalI cleavage sites | This work |
| pBRAXAC2606(1-163) | pBRA vector for the expression of XAC2606(1-163) ,xac2606 orf was inserted at the NcoI-SalI cleavage sites. | This work |
| pBRAVirB5 | pBRA vector for the expression of XAC2613(1-275)(virB5),xac2613 orf was inserted at the NcoI-SalI cleavage sites. | This work |
| pBRA-SmIt2994-2995-2996 | pBRA vector expression of SmIt2994, SmIt2995, SmIt 2996 from *S. maltophilia* K279a | This work |
| pBRA-SmIt2994-5-6-7-8 | pBRA vector expression of SmIt2994, SmIt2995, SmIt 2996, SmIt2997(virB6), SmIt2998(VirB5) from *S. maltophilia* K279a | This work |
| pBRA-SmIt2994 | pBRA vector for the expression of SmIt2994 | This work |
| pBRA-SmIt2995 | pBRA vector for the expression of SmIt2995 | This work |
| pBRA-SmIt2996 | pBRA vector for the expression of SmIt2996 | This work |
| pNPTS138-ΔXAC2611 | Suicide vector for deletion of xac2611 in the *X. citri* Δxac2611, Δ8Δ2609ΔXAC2611, Δ8Δ2609ΔXAC2611ΔXAC2606. Km<sup>r</sup> | This work |
| pNPTS138-Δxac2606 | Suicide vector for deletion of xac2606 in the *X. citri* Δxac2606, Δ8Δ2609ΔXAC2611ΔXAC2606. Km<sup>r</sup> | This work |
| pNPTS138-Δxac2613(ΔvirB5) | Suicide vector for deletion of xac2613(virB5) in the *X. citri* Δxac2613, Δxac2613Δxac2611. | Oka et al, 2024 |
| pNPTS138--ΔXAC209--ΔXAC2610-ΔXAC2611 | Suicide vector for deletion of locus xac2609-xac2610- xac2611 in the *X. citri* ΔX-Tfe2609-X-Tfi2610-Δxac2611. | This work |
| William plasmid |  |  |

**S3 Table.**
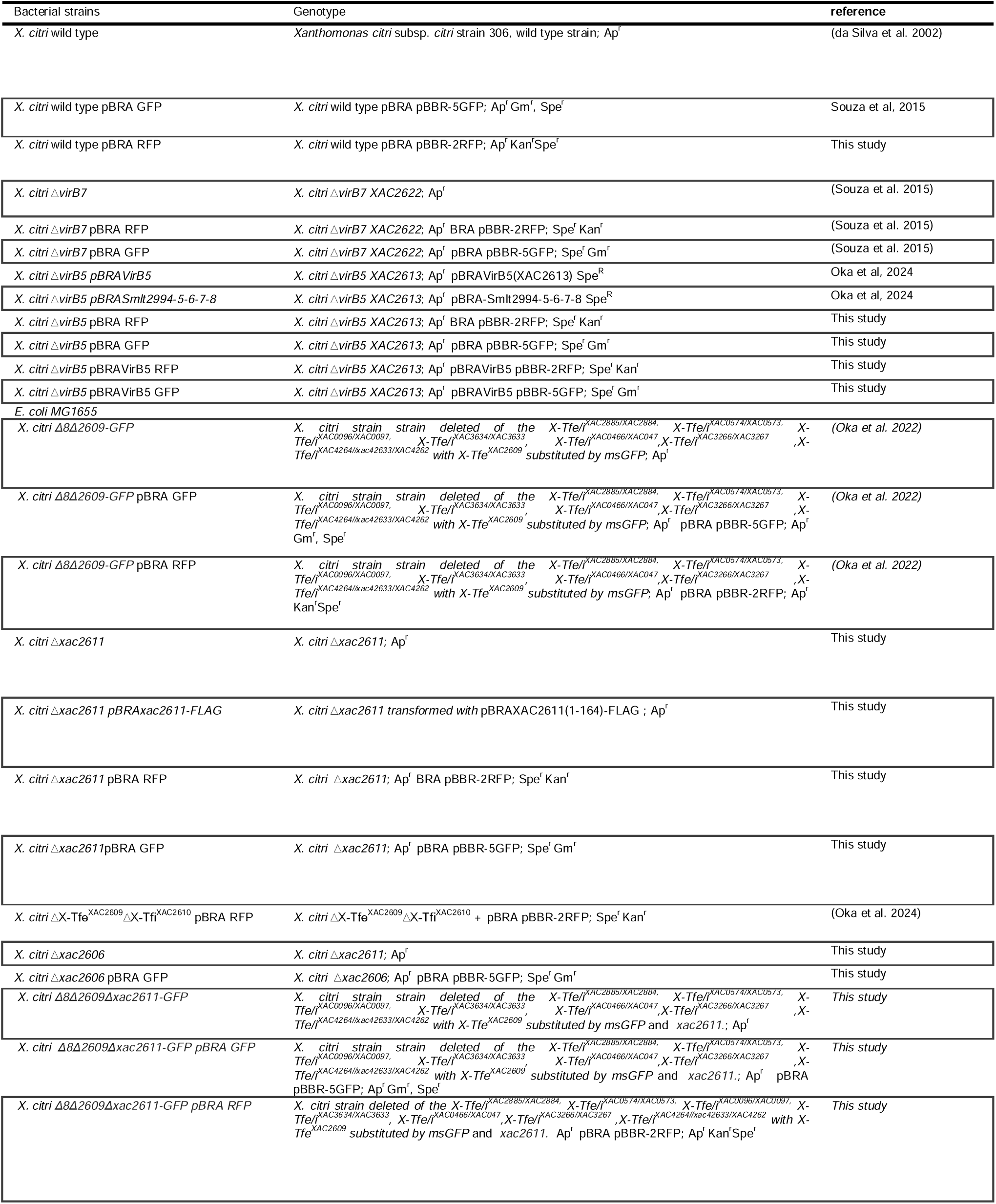
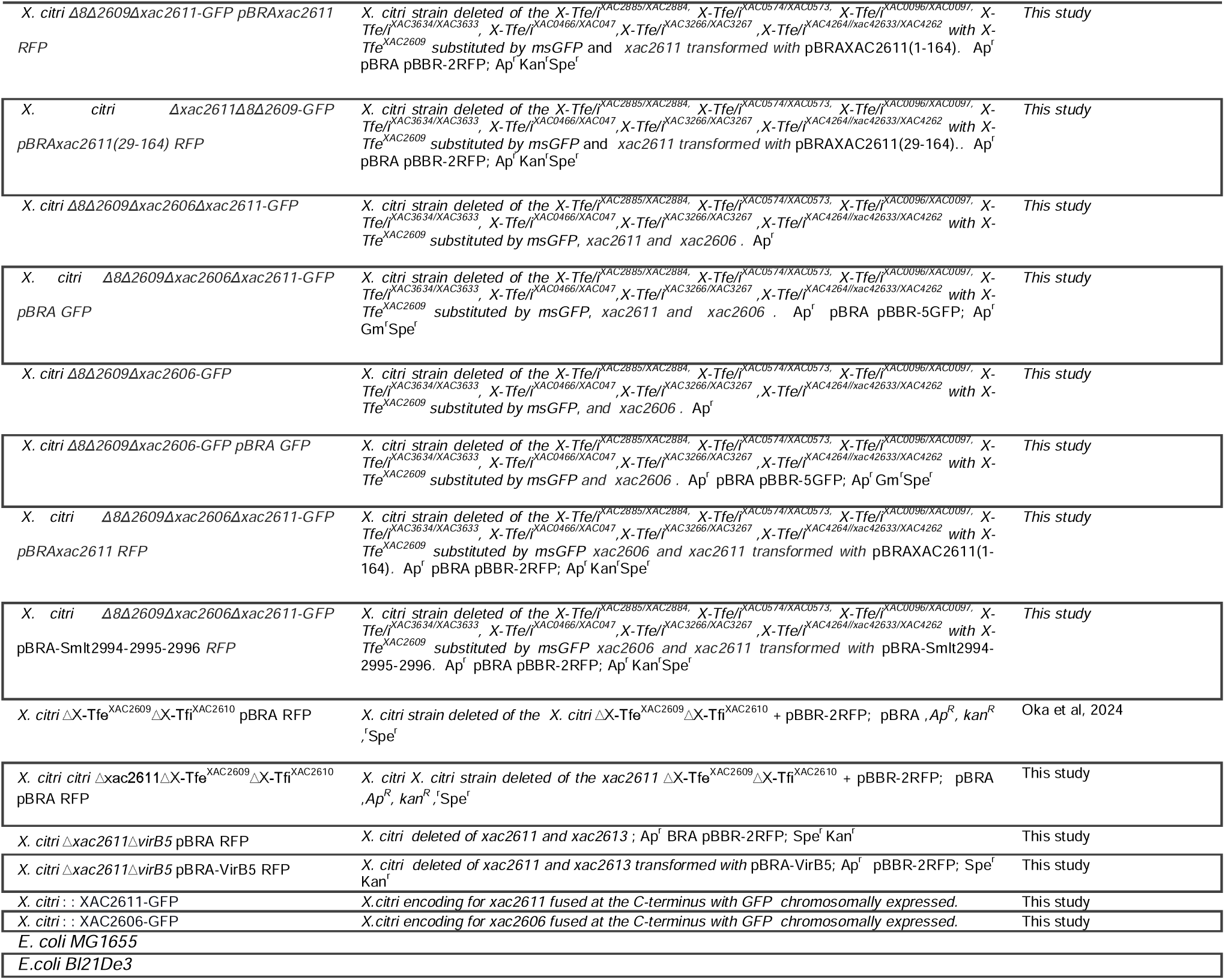
Bacterial strains used in this study.

### Supplementary Material S1. Tukey HSD significance matrix for **Figure 5**)

This supplementary file contains the complete Tukey HSD significance matrix corresponding to the statistical analysis shown in **Figure 5**). Pairwise comparisons were performed on log10-transformed gentamicin-to-kanamycin CFU ratios obtained from independent *X. citri* co-culture competition assays. The matrix reports whether each pairwise comparison was statistically significant after Tukey HSD multiple-comparison correction. “ns” indicates non-significant comparisons with adjusted *p* ≥ 0.05, whereas “*” indicates significant comparisons with adjusted p < 0.05. Diagonal entries correspond to self-comparisons of each condition. The WT GFP versus WT RFP condition was used as the reference control for significance annotation in the main* **Figure 5**).

### Supplementary Movies Legend

- **Supplementary Movie 1** – *Wild-type X. citri (RFP)* monoculture control. Cells maintain morphology and red fluorescence throughout 5-hour time-lapse, indicating normal growth without lysis. Images were acquired at 10-minute intervals and show merged micrographs of brightfield and red fluorescence channels from cells grown on LB-agarose media. Scale bar 5 um.
- **Supplementary Movie 2** – Δ*xac2611 (RFP)* monoculture. Cells lacking XAC2611 display loss of fluorescence and morphological collapse, indicating extensive X-T4SS-mediated lysis by sibling cells. Images were acquired at 10-minute intervals and show merged micrographs of brightfield and red fluorescence channels from cells grown on LB-agarose media. Scale bar 5 um.
- **Supplementary Movie 3** – Δ*xac2611*Δ*virB5 (RFP)* monoculture. Double mutant lacking both XAC2611 and VirB5 does not undergo lysis, indicating that X-T4SS is required for the observed cell death. Images were acquired at 10-minute intervals and show merged micrographs of brightfield and red fluorescence channels from cells grown on LB-agarose media. Scale bar 5 um.
- **Supplementary Movie 4** – Δ*xac2611*Δ*virB5 (RFP)* complemented with plasmid expressing VirB5. Lysis is restored, confirming that VirB5-mediated X-T4SS activity causes trans-intoxication in absence of XAC2611.Images were acquired at 10-minute intervals and show merged micrographs of brightfield and red fluorescence channels from cells grown on LB-agarose media. Scale bar 5 um.
- **Supplementary Movie 5** – Δ*8*Δ*2609*Δ*xac2611 (GFP)* strain (lacking all major X-Tfes) shows no cell death despite lacking XAC2611, demonstrating that effectors are required for trans-intoxication.Images were acquired at 10-minute intervals and show merged micrographs of brightfield and green fluorescence channels from cells grown on LB-agarose media. Scale bar 5 um.
- **Supplementary Movie 6** – Δ*8*Δ*2609*Δ*xac2611 + pBRA-XAC2611* strain. Complementation with XAC2611 restores protection and eliminates cell death, confirming the protective role of XAC2611. Images were acquired at 10-minute intervals and show brightfield micrographs from cells grown on LB-agarose media. Scale bar 5 um.
- **Supplementary Movie 7** – Co-culture: *WT-GFP vs* Δ*8*Δ*2609-RFP*. No lysis of Δ8Δ2609 strain, confirming that X-Tfis are not essential to prevent fratricide in this background. Images were acquired at 10-minute intervals and display merged micrographs of brightfield, red, and green fluorescence channels from cells grown on LB-agarose media. Scale bar: 5 µm. Objective: 100×.
- **Supplementary Movie 8** – Co-culture: *WT-GFP vs* Δ*8*Δ*2609*Δ*xac2611-RFP*. Strong lysis of recipient cells demonstrates that XAC2611 is required to prevent trans-intoxication by wild-type siblings. Images were acquired at 10-minute intervals and display merged micrographs of brightfield, red, and green fluorescence channels from cells grown on LB-agarose media. Scale bar: 5 µm. Objective: 100×.
- **Supplementary Movie 9** – Co-culture: Δ*8*Δ*2609-GFP vs* Δ*8*Δ*2609*Δ*xac2611-RFP*. No lysis observed, confirming that killing requires both a functional T4SS and presence of effectors in donor cells. Images were acquired at 10-minute intervals and display merged micrographs of brightfield, red, and green fluorescence channels from cells grown on LB-agarose media. Scale bar: 5 µm. Objective: 100×.
- **Supplementary Movie 10** – Co-culture: *WT-GFP vs* Δ*8*Δ*2609*Δ*xac2606-RFP*. No significant killing was detected, indicating that XAC2606 is not the key factor for fratricide protection. Images were acquired at 10-minute intervals and display merged micrographs of brightfield, red, and green fluorescence channels from cells grown on LB-agarose media. Scale bar: 5 µm. Objective: 100×.
- **Supplementary Movie 11** – Co-culture: Δ*virB5-GFP vs* Δ*8*Δ*2609*Δ*xac2611-RFP*. No lysis observed due to absence of VirB5 in donor, confirming that VirB5 is essential for T4SS-mediated killing. Images were acquired at 10-minute intervals and display merged micrographs of brightfield, red, and green fluorescence channels from cells grown on LB-agarose media. Scale bar: 5 µm. Objective: 100×.
- **Supplementary Movie 12** – Co-culture: Δ*virB5 + pBRA-VirB5-GFP vs* Δ*8*Δ*2609*Δ*xac2611-RFP*. Lysis is restored with VirB5 complementation, confirming its role in pilus-mediated effector delivery.Images were acquired at 10-minute intervals and display merged micrographs of brightfield, red, and green fluorescence channels from cells grown on LB-agarose media. Scale bar: 5 µm. Objective: 100×.
- **Supplementary Movie 13** – Co-culture: *WT-GFP vs* Δ*8*Δ*2609*Δ*xac2611 + pBRA-XAC2611-RFP*. Complementation of XAC2611 in recipient restores protection and abrogates cell death.Images were acquired at 10-minute intervals and display merged micrographs of brightfield, red, and green fluorescence channels from cells grown on LB-agarose media. Scale bar: 5 µm. Objective: 100×.\
- **Supplementary Movie 14. Coculture of *S. maltophilia* WT with *X. citri* GFP** Δ**8**Δ**2609**Δ***xac*2606**Δ**xac2611 + pBRA-empty.** Four representative microscopic fields are shown. Each field was monitored for 8 h by time-lapse microscopy at 10 min intervals on AB medium microscopy pads at room temperature. Merged channels are shown, with brightfield in gray and GFP fluorescence in green. Scale bar, 5 µm. Objective, 100×.
- **Supplementary Movie 15. Coculture of *S. maltophilia* WT with *X. citri* GFP** Δ**8**Δ**2609**Δ***xac*2606**Δ**xac2611 + pBRA-Smlt2994.** Four representative microscopic fields are shown. Each field was monitored for 8 h by time-lapse microscopy at 10 min intervals on AB medium microscopy pads at room temperature. Merged channels are shown, with brightfield in gray and GFP fluorescence in green. Scale bar, 5 µm. Objective, 100×.
- **Supplementary Movie 16. Coculture of *S. maltophilia* WT with *X. citri* GFP** Δ**8**Δ**2609**Δ***xac*2606**Δ**xac2611 + pBRA-Smlt2995.** Four representative microscopic fields are shown. Each field was monitored for 8 h by time-lapse microscopy at 10 min intervals on AB medium microscopy pads at room temperature. Merged channels are shown, with brightfield in gray and GFP fluorescence in green. Scale bar, 5 µm. Objective, 100×.
- **Supplementary Movie 17. Coculture of *S. maltophilia* WT with *X. citri* GFP** Δ**8**Δ**2609**Δ**xac2611 + pBRA-Smlt2996.** Four representative microscopic fields are shown. Each field was monitored for 8 h by time-lapse microscopy at 10 min intervals on AB medium microscopy pads at room temperature. Merged channels are shown, with brightfield in gray and GFP fluorescence in green. Scale bar, 5 µm. Objective, 100×.

