## Supplementary figures and images for "To Kill or not to Kill: A Conserved *trans*-intoxication protection factor Blocks X-T4SS-Mediated Fratricide through Interaction with VirB5"

### Supplementary Figure 1

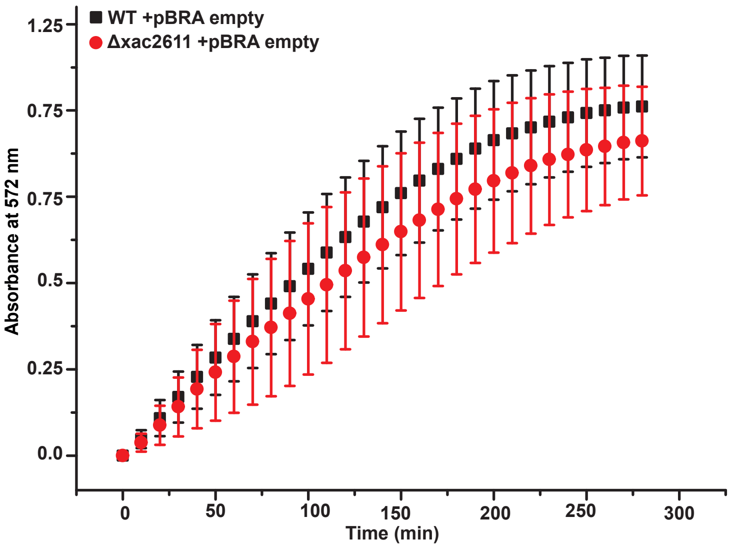

### Supplementary Figure 2

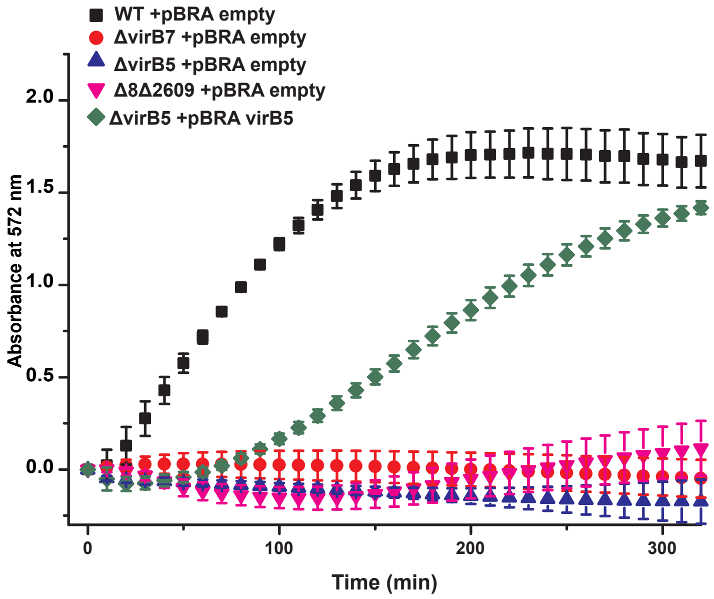
