## Supplementary Material S1 for "To Kill or not to Kill: A Conserved *trans*-intoxication protection factor Blocks X-T4SS-Mediated Fratricide through Interaction with VirB5"

### Tukey adjusted p-value matrix

| | WT GFP vs WT RFP | WT GFP vs ΔvirB7RFP | WT GFP vs ΔvirB5RFP | WT GFP vs Δ8Δ2609RFP | WT GFP vs Δ8Δ2609ΔXAC2611RFP | WT GFP vs Δ8Δ2609ΔXAC2606RFP | WT GFP vs Δ8Δ2609ΔXAC2611RFP + XAC2611 | WT GFP vs Δ8Δ2609ΔXAC2606ΔXAC2611RFP | WT GFP vs Δ8Δ2609ΔXAC2606ΔXAC2611RFP + XAC2611 | WT GFP vs ΔXAC2611RFP | WT GFP vs ΔXAC2610ΔXAC2609 RFP | WT GFP vs Δ8Δ2609ΔXAC2606ΔXAC2611 RFP + Smlt2994-5-6 | WT GFP vs Δ8Δ2609ΔXAC2611 RFP + XAC2611cyt | ΔvirB7 GFP vs WT RFP | ΔvirB7 GFP vs Δ8Δ2609RFP | ΔvirB7 GFP vs ΔXAC2611RFP | ΔvirB7 GFP vs Δ8Δ2609ΔXAC2611RFP | ΔvirB7 GFP vs Δ8Δ2609ΔXAC2606ΔXAC2611RFP | ΔvirB7 GFP vs Δ8Δ2609ΔXAC2606RFP | ΔvirB7 GFP vs Δ8Δ2609ΔXAC2611RFP + XAC2611 | ΔvirB7 GFP vs ΔvirB7RFP | ΔvirB7 GFP vs Δ8Δ2609ΔXAC2611RFP + XAC2611cyt | Δ8Δ2609 GFP vs WT RFP | Δ8Δ2609 GFP vs Δ8Δ2609ΔXAC2611RFP | Δ8Δ2609ΔXAC2611GFP vs WT RFP | Δ8Δ2609ΔXAC2611GFP vs ΔvirB7 RFP | Δ8Δ2609ΔXAC2611GFP vs ΔvirB5 RFP | Δ8Δ2609ΔXAC2611GFP vs Δ8Δ2609 RFP |
|---|---|---|---|---|---|---|---|---|---|---|---|---|---|---|---|---|---|---|---|---|---|---|---|---|---|---|---|---|
| WT GFP vs WT RF | 1.0000 | 0.9969 | 1.0000 | 1.0000 | <0.0001 | 0.0002 | 0.1195 | <0.0001 | <0.0001 | <0.0001 | 1.0000 | <0.0001 | <0.0001 | 1.0000 | 1.0000 | 0.1201 | 1.0000 | 0.9984 | 1.0000 | 1.0000 | 1.0000 | 1.0000 | 1.0000 | 1.0000 | <0.0001 | 1.0000 | 0.9967 | 1.0000 |
| WT GFP vs ΔvirB7RF | 0.9969 | 1.0000 | 1.0000 | 1.0000 | <0.0001 | 0.1037 | 0.9789 | <0.0001 | 0.0050 | <0.0001 | 1.0000 | <0.0001 | <0.0001 | 0.8008 | 1.0000 | 0.9158 | 0.9990 | 1.0000 | 1.0000 | 0.9999 | 0.9565 | 1.0000 | 0.7707 | 1.0000 | <0.0001 | 0.9717 | 0.5029 | 0.9520 |
| WT GFP vs ΔvirB5RF | 1.0000 | 1.0000 | 1.0000 | 1.0000 | <0.0001 | 0.1370 | 0.9557 | <0.0001 | 0.0116 | <0.0001 | 1.0000 | <0.0001 | <0.0001 | 0.9969 | 1.0000 | 0.8660 | 1.0000 | 1.0000 | 1.0000 | 1.0000 | 0.9998 | 1.0000 | 0.9883 | 1.0000 | <0.0001 | 0.9997 | 0.8974 | 0.9991 |
| WT GFP vs Δ8Δ2609R | 1.0000 | 1.0000 | 1.0000 | 1.0000 | <0.0001 | 0.0177 | 0.7242 | <0.0001 | 0.0006 | <0.0001 | 1.0000 | <0.0001 | <0.0001 | 0.9985 | 1.0000 | 0.5942 | 1.0000 | 1.0000 | 1.0000 | 1.0000 | 1.0000 | 1.0000 | 0.9932 | 1.0000 | <0.0001 | 0.9999 | 0.9163 | 0.9997 |
| FP vs Δ8Δ2609ΔXAC2 | <0.0001 | <0.0001 | <0.0001 | <0.0001 | 1.0000 | <0.0001 | <0.0001 | 1.0000 | <0.0001 | <0.0001 | <0.0001 | 1.0000 | 0.0296 | <0.0001 | <0.0001 | <0.0001 | <0.0001 | <0.0001 | <0.0001 | <0.0001 | <0.0001 | <0.0001 | <0.0001 | <0.0001 | <0.0001 | <0.0001 | <0.0001 | <0.0001 |
| FP vs Δ8Δ2609ΔXAC2 | 0.0002 | 0.1037 | 0.1370 | 0.0177 | <0.0001 | 1.0000 | 0.9997 | <0.0001 | 1.0000 | <0.0001 | 0.0177 | <0.0001 | <0.0001 | <0.0001 | 0.0032 | 1.0000 | 0.0006 | 0.4551 | 0.0473 | 0.0037 | 0.0003 | 0.0476 | 0.0001 | 0.0304 | <0.0001 | 0.0022 | <0.0001 | 0.0015 |
| Δ8Δ2609ΔXAC2611R | 0.1195 | 0.9789 | 0.9557 | 0.7242 | <0.0001 | 0.9997 | 1.0000 | <0.0001 | 0.8705 | <0.0001 | 0.5948 | <0.0001 | <0.0001 | 0.0193 | 0.4306 | 1.0000 | 0.1906 | 0.9994 | 0.8030 | 0.3548 | 0.0788 | 0.7204 | 0.0312 | 0.7127 | <0.0001 | 0.1616 | 0.0144 | 0.1286 |
| Δ8Δ2609ΔXAC2606Δ | <0.0001 | <0.0001 | <0.0001 | <0.0001 | 1.0000 | <0.0001 | <0.0001 | 1.0000 | <0.0001 | <0.0001 | <0.0001 | 1.0000 | 0.1271 | <0.0001 | <0.0001 | <0.0001 | <0.0001 | <0.0001 | <0.0001 | <0.0001 | <0.0001 | <0.0001 | <0.0001 | <0.0001 | <0.0001 | <0.0001 | <0.0001 | <0.0001 |
| 609ΔXAC2606ΔXAC2 | <0.0001 | 0.0050 | 0.0116 | 0.0006 | <0.0001 | 1.0000 | 0.8705 | <0.0001 | 1.0000 | 0.0020 | 0.0009 | <0.0001 | <0.0001 | <0.0001 | <0.0001 | 0.9998 | <0.0001 | 0.0715 | 0.0029 | 0.0001 | <0.0001 | 0.0038 | <0.0001 | 0.0017 | <0.0001 | 0.0001 | <0.0001 | <0.0001 |
| WT GFP vs ΔXAC2611 | <0.0001 | <0.0001 | <0.0001 | <0.0001 | <0.0001 | <0.0001 | <0.0001 | <0.0001 | 0.0020 | 1.0000 | <0.0001 | <0.0001 | 0.7533 | <0.0001 | <0.0001 | 0.0002 | <0.0001 | <0.0001 | <0.0001 | <0.0001 | <0.0001 | <0.0001 | <0.0001 | <0.0001 | <0.0001 | <0.0001 | <0.0001 | <0.0001 |
| P vs ΔXAC2610ΔXAC | 1.0000 | 1.0000 | 1.0000 | 1.0000 | <0.0001 | 0.0177 | 0.5948 | <0.0001 | 0.0009 | <0.0001 | 1.0000 | <0.0001 | <0.0001 | 1.0000 | 1.0000 | 0.4655 | 1.0000 | 1.0000 | 1.0000 | 1.0000 | 1.0000 | 1.0000 | 1.0000 | 1.0000 | <0.0001 | 1.0000 | 0.9973 | 1.0000 |
| 9ΔXAC2606ΔXAC261 | <0.0001 | <0.0001 | <0.0001 | <0.0001 | 1.0000 | <0.0001 | <0.0001 | 1.0000 | <0.0001 | <0.0001 | <0.0001 | 1.0000 | 0.1906 | <0.0001 | <0.0001 | <0.0001 | <0.0001 | <0.0001 | <0.0001 | <0.0001 | <0.0001 | <0.0001 | <0.0001 | <0.0001 | <0.0001 | <0.0001 | <0.0001 | <0.0001 |
| 8Δ2609ΔXAC2611 RF | <0.0001 | <0.0001 | <0.0001 | <0.0001 | 0.0296 | <0.0001 | <0.0001 | 0.1271 | <0.0001 | 0.7533 | <0.0001 | 0.1906 | 1.0000 | <0.0001 | <0.0001 | <0.0001 | <0.0001 | <0.0001 | <0.0001 | <0.0001 | <0.0001 | <0.0001 | <0.0001 | <0.0001 | <0.0001 | <0.0001 | <0.0001 | <0.0001 |
| ΔvirB7 GFP vs WT R | 1.0000 | 0.8008 | 0.9969 | 0.9985 | <0.0001 | <0.0001 | 0.0193 | <0.0001 | <0.0001 | <0.0001 | 1.0000 | <0.0001 | <0.0001 | 1.0000 | 0.9999 | 0.0245 | 1.0000 | 0.8882 | 1.0000 | 1.0000 | 1.0000 | 1.0000 | 1.0000 | 1.0000 | <0.0001 | 1.0000 | 1.0000 | 1.0000 |
| irB7 GFP vs Δ8Δ260 | 1.0000 | 1.0000 | 1.0000 | 1.0000 | <0.0001 | 0.0032 | 0.4306 | <0.0001 | <0.0001 | <0.0001 | 1.0000 | <0.0001 | <0.0001 | 0.9999 | 1.0000 | 0.3541 | 1.0000 | 1.0000 | 1.0000 | 1.0000 | 1.0000 | 1.0000 | 0.9992 | 1.0000 | <0.0001 | 1.0000 | 0.9705 | 1.0000 |
| rB7 GFP vs ΔXAC261 | 0.1201 | 0.9158 | 0.8660 | 0.5942 | <0.0001 | 1.0000 | 1.0000 | <0.0001 | 0.9998 | 0.0002 | 0.4655 | <0.0001 | <0.0001 | 0.0245 | 0.3541 | 1.0000 | 0.1696 | 0.9895 | 0.6624 | 0.2823 | 0.0722 | 0.5721 | 0.0297 | 0.5713 | <0.0001 | 0.1228 | 0.0133 | 0.0989 |
| GFP vs Δ8Δ2609ΔXA | 1.0000 | 0.9990 | 1.0000 | 1.0000 | <0.0001 | 0.0006 | 0.1906 | <0.0001 | <0.0001 | <0.0001 | 1.0000 | <0.0001 | <0.0001 | 1.0000 | 1.0000 | 0.1696 | 1.0000 | 0.9993 | 1.0000 | 1.0000 | 1.0000 | 1.0000 | 1.0000 | 1.0000 | <0.0001 | 1.0000 | 0.9977 | 1.0000 |
| s Δ8Δ2609ΔXAC260 | 0.9984 | 1.0000 | 1.0000 | 1.0000 | <0.0001 | 0.4551 | 0.9994 | <0.0001 | 0.0715 | <0.0001 | 1.0000 | <0.0001 | <0.0001 | 0.8882 | 1.0000 | 0.9895 | 0.9993 | 1.0000 | 1.0000 | 0.9999 | 0.9726 | 1.0000 | 0.8378 | 1.0000 | <0.0001 | 0.9770 | 0.5927 | 0.9619 |
| GFP vs Δ8Δ2609ΔXA | 1.0000 | 1.0000 | 1.0000 | 1.0000 | <0.0001 | 0.0473 | 0.8030 | <0.0001 | 0.0029 | <0.0001 | 1.0000 | <0.0001 | <0.0001 | 1.0000 | 1.0000 | 0.6624 | 1.0000 | 1.0000 | 1.0000 | 1.0000 | 1.0000 | 1.0000 | 0.9995 | 1.0000 | <0.0001 | 1.0000 | 0.9803 | 1.0000 |
| s Δ8Δ2609ΔXAC2611 | 1.0000 | 0.9999 | 1.0000 | 1.0000 | <0.0001 | 0.0037 | 0.3548 | <0.0001 | 0.0001 | <0.0001 | 1.0000 | <0.0001 | <0.0001 | 1.0000 | 1.0000 | 0.2823 | 1.0000 | 0.9999 | 1.0000 | 1.0000 | 1.0000 | 1.0000 | 1.0000 | 1.0000 | <0.0001 | 1.0000 | 0.9985 | 1.0000 |
| virB7 GFP vs ΔvirB7 | 1.0000 | 0.9565 | 0.9998 | 1.0000 | <0.0001 | 0.0003 | 0.0788 | <0.0001 | <0.0001 | <0.0001 | 1.0000 | <0.0001 | <0.0001 | 1.0000 | 1.0000 | 0.0722 | 1.0000 | 0.9726 | 1.0000 | 1.0000 | 1.0000 | 1.0000 | 1.0000 | 1.0000 | <0.0001 | 1.0000 | 1.0000 | 1.0000 |
| Δ8Δ2609ΔXAC2611R | 1.0000 | 1.0000 | 1.0000 | 1.0000 | <0.0001 | 0.0476 | 0.7204 | <0.0001 | 0.0038 | <0.0001 | 1.0000 | <0.0001 | <0.0001 | 1.0000 | 1.0000 | 0.5721 | 1.0000 | 1.0000 | 1.0000 | 1.0000 | 1.0000 | 1.0000 | 1.0000 | 1.0000 | <0.0001 | 1.0000 | 0.9994 | 1.0000 |
| 8Δ2609 GFP vs WT F | 1.0000 | 0.7707 | 0.9883 | 0.9932 | <0.0001 | 0.0001 | 0.0312 | <0.0001 | <0.0001 | <0.0001 | 1.0000 | <0.0001 | <0.0001 | 1.0000 | 0.9992 | 0.0297 | 1.0000 | 0.8378 | 0.9995 | 1.0000 | 1.0000 | 1.0000 | 1.0000 | 0.9999 | <0.0001 | 1.0000 | 1.0000 | 1.0000 |
| GFP vs Δ8Δ2609ΔX | 1.0000 | 1.0000 | 1.0000 | 1.0000 | <0.0001 | 0.0304 | 0.7127 | <0.0001 | 0.0017 | <0.0001 | 1.0000 | <0.0001 | <0.0001 | 1.0000 | 1.0000 | 0.5713 | 1.0000 | 1.0000 | 1.0000 | 1.0000 | 1.0000 | 1.0000 | 0.9999 | 1.0000 | <0.0001 | 1.0000 | 0.9915 | 1.0000 |
| 609ΔXAC2611GFP vs | <0.0001 | <0.0001 | <0.0001 | <0.0001 | <0.0001 | <0.0001 | <0.0001 | <0.0001 | <0.0001 | <0.0001 | <0.0001 | <0.0001 | <0.0001 | <0.0001 | <0.0001 | <0.0001 | <0.0001 | <0.0001 | <0.0001 | <0.0001 | <0.0001 | <0.0001 | <0.0001 | <0.0001 | 1.0000 | <0.0001 | <0.0001 | <0.0001 |
| 9ΔXAC2611GFP vs Δ | 1.0000 | 0.9717 | 0.9997 | 0.9999 | <0.0001 | 0.0022 | 0.1616 | <0.0001 | 0.0001 | <0.0001 | 1.0000 | <0.0001 | <0.0001 | 1.0000 | 1.0000 | 0.1228 | 1.0000 | 0.9770 | 1.0000 | 1.0000 | 1.0000 | 1.0000 | 1.0000 | 1.0000 | <0.0001 | 1.0000 | 1.0000 | 1.0000 |
| 9ΔXAC2611GFP vs Δ | 0.9967 | 0.5029 | 0.8974 | 0.9163 | <0.0001 | <0.0001 | 0.0144 | <0.0001 | <0.0001 | <0.0001 | 0.9973 | <0.0001 | <0.0001 | 1.0000 | 0.9705 | 0.0133 | 0.9977 | 0.5927 | 0.9803 | 0.9985 | 1.0000 | 0.9994 | 1.0000 | 0.9915 | <0.0001 | 1.0000 | 1.0000 | 1.0000 |
| ΔXAC2611GFP vs Δ8 | 1.0000 | 0.9520 | 0.9991 | 0.9997 | <0.0001 | 0.0015 | 0.1286 | <0.0001 | <0.0001 | <0.0001 | 1.0000 | <0.0001 | <0.0001 | 1.0000 | 1.0000 | 0.0989 | 1.0000 | 0.9619 | 1.0000 | 1.0000 | 1.0000 | 1.0000 | 1.0000 | 1.0000 | <0.0001 | 1.0000 | 1.0000 | 1.0000 |

Notes: Values are Tukey HSD adjusted p-values. Statistical tests were performed on log10(viability ratio). Cells with adjusted p < 0.05 are highlighted. Control group for main-plot asterisks: WT GFP vs WT RFP.
